# Edge effects constraint endemic but not introduced arthropod species in a pristine forest on Terceira (Azores, Portugal)

**DOI:** 10.1101/2022.09.07.506827

**Authors:** Noelline Tsafack, Gabor Pozsgai, Mario Boieiro, Alejandra Ros-Prieto, Rui Nunes, Maria T Ferreira, Paulo A. V. Borges

## Abstract

Pristine Azorean forests have been deeply fragmented since human colonization. Fragmentation increases the length of edges and it therefore promotes edge habitats. Studying the impact of edge habitat on species assemblages is crucial to highlight the importance of forest connectivity and guide management strategies.

This study explores the impact of forest edges on arthropod assemblages, and particularly investigates the differences of arthropod communities between three habitats, along a distance gradient from the forest edge near a pasture matrix to the core forest. We also compare patterns of arthropod communities with different biogeographic status (endemic, native non-endemic, and introduced species). We sampled in a pristine forest on Terceira island bordered by semi-natural pastures, using flight interception traps. Traps were established along the ecological gradient of three habitats: the forest edge, an intermediate habitat at 100 m from the border, and the core forest at 300 m deep from the border.

We took a multi-taxon approach and used both univariate and multivariate techniques to understand the variation in arthropods species abundance, richness and composition between the three habitats in the native forest.

Overall, endemic species dominated arthropod abundances whereas species richness and diversity were similar between the three biogeographic categories. We found evidence of a strong edge effect on arthropod assemblages, adjusted both by biogeographic categories and seasonality. Indigenous (endemic and native non-endemic) species abundances were higher in the forest interior than at the edges or intermediate habitats, suggesting that indigenous arthropod assemblages were sensitive to the distance from the edge, whereas introduced species abundances did no show an edge effect. Species diversity and richness did not differ between the three habitats either, regardless of the biogeographic categories. The composition of arthropods between the three habitats differed significantly when we considered all species or endemic species only, but not with native non-endemic or introduced species. However, the difference got obscured when seasonality was included in the analyses, suggesting that even though edges impact species composition, this impact varies seasonally and endemic species are particularly affected in early summer.

Our results indicate that forest edges impact arthropods assemblages but endemic species are more likely to be constrained by the increase of edges than introduced species. Since most of these endemic species are of conservation concern, we urge to avoid forest management strategies that increase fragmentation and call for action to increase the size of native forest habitat.

**Highlights:**

- Edge effects constraint the abundance and composition of endemic species but not of introduced ones
- Both biogeographical categories and temporal dynamics play a role in forming assemblage patterns
- Endemics are the most abundant but richness is similar for all three biogeographic categories
- The edge can affect the abundance of indigenous species up to100 m into the forest interior

## 1. Introduction

Island landscapes have been deeply altered since human colonization (Nogué et al., 2017). Particularly native forest fragments have suffered from intensive management including deforestation for wood exploitation, agriculture and urban infrastructures to support the development of human societies in island ecosystems (Braje et al., 2017).

The most important ecological process that resulted from these modifications is native forest fragmentation. Habitat fragmentation involves both habitat loss and the increasing lack of connectivity between habitat remnants leading to significant changes in abiotic and biotic conditions of the remaining patches (Fahrig, 2003). Many studies that investigated the effects of habitat fragmentation on the biota agreed that forest fragmentation poses a substantial threat for biodiversity and ecosystems sustainability (Barbosa and Marquet, 2002; Haila, 2002; Castro et al., 2010; Terraube et al., 2016; Liu et al., 2019; Schlegel, 2022).

A major consequence of habitat fragmentation is the increase of the area of edge habitats. These areas usually show abiotic and biotic affinities with the contiguous habitats, but may also present specific characteristics. Murcia (1995) summarized the main ecological consequences of the edges: abiotic, direct biological and indirect biological effects. (i) Abiotic effects include changes that occur in environmental parameters resulting from the geographical proximity with a different environment. (ii) Direct biological effects include changes in the biota’s community structure (e.g. abundance, richness, or/and composition) caused directly by new environmental conditions at the edges. (iii) Indirect biological effects include changes in species interactions due to the previous abiotic and direct biological effects. Forest edges represent a more or less abrupt transitional zone which may support species communities well adapted to this type of environment or might serve as a secondary refuge for species from adjacent habitats following a source-sink dynamic. In the rare studies that investigated edge effects in island forest habitats, the consensus is that edge habitats are detrimental for species diversity and abundance (Otto et al., 2014). However, this might depend on the composition of the assemblage. For example, introduced (exotic) species are known to be more resilient to environment changes while indigenous species (endemic and native species) are more sensitive (Borges et al., 2020).

In this study, we aimed to test the edge effects in a native forest fragment on Terceira island where the most important areas of Azorean native forests have been preserved (Gaspar et al., 2008, 2011). We compared arthropod communities of three habitats in a distance gradient from the edge habitat to the core forest. We also compared patterns of arthropod communities with different biogeographic status (endemic, native non-endemic, and introduced species) and tested if sampling season influences edge effects.

We addressed the following hypotheses:

i. The abundance and richness in indigenous species (endemic and native non-endemic species) community will be increasing from the edge to inside the forest. Dominant species, however, will spread from the deep forest toward the edges.
ii. Exotic species will show opposite patterns and will be more abundant and diverse in edge habitat due to proximity of semi-natural pastures.
iii. The assemblage composition of the three habitats will be different but forest edge will markedly be different from both center and deep forest.
iv. There will be seasonal differences in how well the assemblages in the three habitats are separated, with the expectation that marked differences will be shown in the most productive seasons (Spring and Summer).

To test these hypotheses, we used a diversity-profile approach based on Hill numbers. Hill numbers allow a complete characterization of the diversity of a community combining information on species richness, species rarity, and species dominance (Chao et al., 2014; Tsafack et al., 2021). We also investigated differences in community composition, using multivariate methods, and finally, we tested if the differences in community species abundances, richness, diversity and composition show seasonal variability.

## 2. Materials and methods

### 2.1. Study area

The study was carried out in Terceira, one of the nine islands of the Azores archipelago (Portugal). Terceira belongs to the Central Group of islands, it is the third largest (402 km^2^) island of the archipelago (Fig. 1 A) and is roughly circular.

**Fig. 1.**
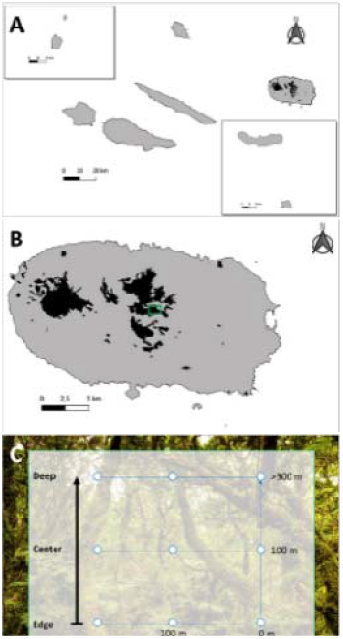
Study area and sampling design: (A) Terceira is located in the Azorean archipelago; (B) Terceira native forest fragments are represented and the Terra-Brava native forest fragment with the experiment area in the center of Terceira is highlighted with green; (C) sampling design with location of SLAM traps from border to the forest center.

Terceira has the largest area of native forest in the archipelago and some of the most pristine forest fragments (Gaspar et al., 2008, 2011). The native forest had covered most of the land surface before human colonization, but suffered a drastic reduction of over 90% during the past five centuries (Fernández-Palacios et al., 2011; Gaspar et al., 2008; Triantis et al., 2010). In Terceira, only five native forest fragments survived the severe human impact, occupying currently ∼6% of the island area (i.e. 23 km^2^) and being mostly restricted to elevations above 500 m a.s.l. The dominant tree and shrub species in the Azorean native forests are the endemic *Erica azorica* Hochst. ex Seub., *Ilex azorica* Gand, *Juniperus brevifolia* (Seub.) Antoine, *Laurus azorica* (Seub.) Franco and *Vaccinium cylindraceum* Sm (Elias et al., 2016; Rego et al., 2019). These native forest fragments harbor a high diversity of endemic arthropods and were included in the recently created Terceira Island Natural Park (Borges et al., 2005; Fattorini et al., 2012).

The study was conducted in the Terra Brava native forest fragment (38º 43’ 56.4” N, 27º 11’ 52.7” W) (Fig. 1B). This fragment of native forest was classified as “Juniperus-Ilex montane forests” being dominated by the endemic trees *J. brevifolia* and *I. azorica*, but with *L. azorica* also being present in high densities (Elias et al., 2016). The dense cover of bryophytes and ferns in all substrates is a typical characteristic of the forest (Gabriel and Bates, 2005). Terra Brava is one of the most pristine native forest fragments (Gaspar et al., 2008, 2011), it is 1.8 km^2^ in area, and is surrounded by a matrix dominated by semi-natural pastures that are mostly used by cattle in the spring and summer.

### 2.2. Experimental design and data collection

We used SLAM (Sea, Land, and Air Malaise) traps to sample the terrestrial arthropod communities in Terra-Brava. SLAM is a type of passive flight interception trap suitable to sample mobile arthropods. This trap is a modified version of the traditional Malaise trap, 110×110×110 cm dimensions, and it consists of a frame of shock-corded poles to which the netting trap clips, allowing the interception of arthropods moving from four different directions. A central white mesh funnels the individuals to a sampling bottle at the top of the trap. sampling bottles contained propylene glycol as a preservative and the traps were monitored every month, from June 2014 to December 2015. The sampled specimens were transported to the lab, sorted and identified to species or morphospecies level (Borges et al., 2022).

We set three linear transects along the ecological gradient that extends from the edge to the core of the native forest fragment (Fig. 1 C). Three SLAM traps were set in each transect: one at the border of the forest fragment (EDGE), a second distanced 100 m from edge (CENTER) and a third at approximately 300 m from edge (DEEP) in the inner area of the forest fragment. Overall, nine SLAM traps were placed in the native forest fragment of Terra-Brava (Fig. 1 C). The forest edge was easy to identify due to the marked differences in plant species composition and structure with neighboring matrix habitat (semi-natural pastures).

We adopted a multi-taxon approach considering most groups of terrestrial arthropods sampled with SLAM traps (Arachnida, Chilopoda, Diplopoda, Insecta (excluding Diptera, Hymenoptera and Lepidoptera)). The different arthropods groups play key ecological roles in forest ecosystems (e.g., herbivores, detritivores, predators) and their diversity, taxonomy and ecology are well known in the Azores (Borges et al., 2010).

Arthropod species were classified into three biogeographical categories according to their colonization status following the Azorean biodiversity checklist (Borges et al., 2010). We considered a species as endemic when its distribution is restricted to the Azores, native non-endemic those species that arrived naturally to the archipelago but are also present elsewhere, and as introduced species those that have been introduced (accidentally or deliberately) by humans in the Azores. The complete dataset is available in GBIF (Borges and Lamelas-López, 2022) and described in detail in Borges et al. (2022).

### 2.3. Data analysis

For the comprehensive understanding of the variation in arthropods species richness, composition and abundance between the three habitats, we conducted complementary statistical analyses using both univariate and multivariate techniques.

All analyses were conducted on the complete data set (with all species) as well as on data subsets which included species sharing the same biogeographical category (i.e. endemic, native non-endemic and introduced species). Overall, we used four species-site matrices populated with count data for the different analysis.

#### Species richness, diversity and abundance

We assessed the diversity patterns in the three study habitats and modeled the responses of abundance and diversity indices to habitat and sampling month (and the interaction of these two variables).

We calculated species diversity indices (Chao species richness, Shannon diversity and Simpson diversity indices) following the Hills numbers as suggested by Chao et al. (2014). Hill numbers are based on the same formula and differ only by an exponent, the parameter *q*. The parameter *q* determines the sensitivity to species relative abundances (Chao et al., 2014). We used the first three hill numbers to express respectively species richness (*q = 0*), Shannon diversity (*q = 1*) and Simpson diversity (*q = 2*). These three numbers give emphasis respectively on rare, common and dominant species (Chao et al., 2014). We calculated the Hill numbers with the *iNEXT* function in the R package iNEXT (Hsieh et al., 2016). We used linear mixed-effects models to test the effect of habitat and sampling period (month) on arthropods abundance and diversity. Analysis of Variance (ANOVA) was used to assess significance of habitat effect followed by a pairwise comparison Tukey test to evaluate the difference between habitat pairs. Habitat was set as a fixed factor and the sites inside habitats as random factors in the ANOVA model.

We also used generalized linear mixed models (GLMMs) to assess the effects of sampling period (month), habitat and their interaction on species abundance and diversity indices.

We used function *lme* in R package nlme for linear mixed-effect models; functions *anova* and *glht* in R package multcomp respectively for ANOVA and for Tukey tests; function *glmmTMB* in glmmTMB R package (Brooks et al., 2017) for GLMM, function *r*.*squaredGLMM* in the R package MuMIn (Bartoń, 2020) to calculate the variance explained (R2) by fixed and random factors. Package ggplot2 (Wickham, 2016) was used for all graphics. All analyses were conducted with the statistical software R (R Core Team, 2021).

#### Species composition and association with the study habitats

Count data were transformed using the Hellinger transformation implemented in the vegan R package (Oksanen et al., 2010), and the arthropod assemblages were compared based on Euclidean distances (O’Hara and Kotze, 2010).

To fully understand the temporal aspects of the sampling regime, and to investigate the effect of the three habitats and the seasonality on arthropods assemblages, two different distance-based redundancy analyses (dbRDA) were carried out on each data matrices.

In the first approach, the effects of sampling year and month were partialled out by setting the ‘Condition’ term to the combination of these two variables (i.e. setting them as random effects) in a partial dbRDA.

In the second approach, along with habitat, the sampling year and sampling month were also included in the model as interacting explanatory variables (i.e. as fixed effects). A permutation test with 999 permutations was used to assess the significance of the model, and the significance of the constraining variable (habitat or habitat × sampling year × sampling month) was estimated by using an ANOVA-like permutation test (with the function *anova*.*cca*, implemented in vegan) with 999 permutations.

The resulting eight dbRDA models were compared based on their Akaike Information Criterion (AIC) values and the variance they explained. Sampling points were plotted based on their dbRDA scores and convex hulls (embracing all points) and ellipses in one standard deviation distance from the centroid were drawn for visual investigation.

We used non-metric multidimensional scaling (NMDS) to ordinate and visualize the differences in arthropod assemblage compositions across sampling months and habitats. NMDS was conducted on the Hellinger-transformed species-site data and the centroid of each sampling month-habitat point groups was calculated and plotted against the first two NMDS axes.

Pairwise Euclidean distances were calculated in each month between samples from different habitats (between-habitat distances) as well as between those from the same habitat (within-habitat distances), while temporal patterns were investigated visually.

## 3. Results

### 3.1. Overall patterns

Overall, we collected a total of 13.516 individuals belonging to 107 (morpho)species. Endemic species was the most abundant group (60%) followed by native non-endemic species (36%) and introduced species only accounted for 4% of the individuals. Species richness was balanced between the three biogeographical categories: we found that 26% of species were endemic, 35% were native non-endemic and 39% were introduced (Fig. 2, Table. A. 1).

**Fig. 2.**
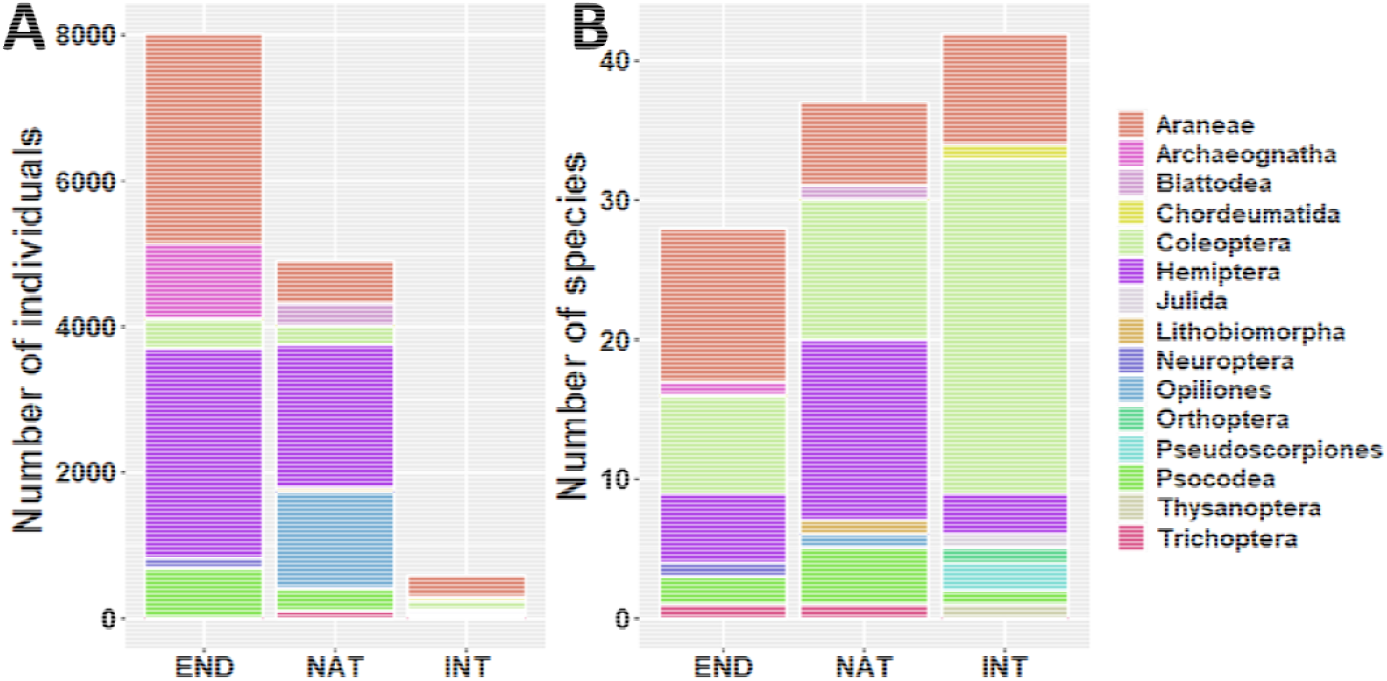
Arthropods orders sampled with SLAM in the three habitats (EDGE, CENTER, DEEP) classified within their biogeographical categories (END - endemic species, NAT - native non-endemic species and INT - introduced species). Overall number of individuals (A) and number of species (B). The graphic should be displayed in color to differentiate the fifteen orders.

Hemiptera was the most abundant order with 4877 individuals (representing 36% of the total) followed by Araneae with 3786 individuals (28%) and Opiliones with 1335 individuals (10%). Only 760 individuals (6%) belonged to Coleoptera (Fig. A.1, Table. A. 1).

Twenty species represented 90% of the total abundance and the four most abundant species were the endemic *Cixius azoterceirae* (Hemiptera, 17%), the native non-endemic *Leiobunum blackwalli* (Opiliones, 10%), the endemic species *Rugathodes acoreensis* (Araneae, 8%) and the endemic *Trigoniophthalmus borgesi* (Archaeognatha, 8%) accounting for about 50 % of the total abundance

Despite being low in abundance, Coleoptera was the most speciose order with 41 species (38%) followed by Araneae with 25 species (23%) and Hemiptera with 21 species (20%).

### 3.2. Edge effect on species abundance, richness and diversity

#### Abundance

Overall, arthropod abundance was substantially higher in the DEEP forest habitat than in the CENTER and EDGE habitats. We observed the same pattern when we considered endemic and native non-endemic species groups, but we found no difference in the introduced species abundances between the three habitats (Fig. 3, Table A.2).

**Fig. 3.**
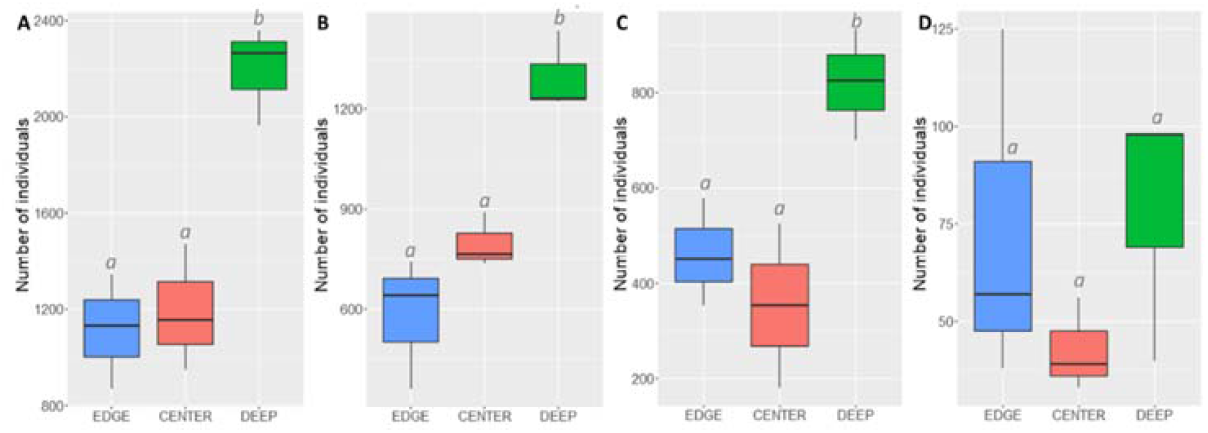
Arthropod abundance in three habitats (forest EDGE, CENTER and DEEP) of Terra-Brava native forest fragment. Boxplots present abundance data for all species A), and for the three biogeographic groups: endemic B), native non-endemic, C) and introduced species D).

#### Species richness and diversity

We found no difference in species richness or diversity metrics between the three habitats, regardless of whether we considered the Chao species richness (Fig. 4A), Shannon diversity (Fig. 5A) or Simpson diversity (Fig. 6A) indexes.

**Fig. 4.**
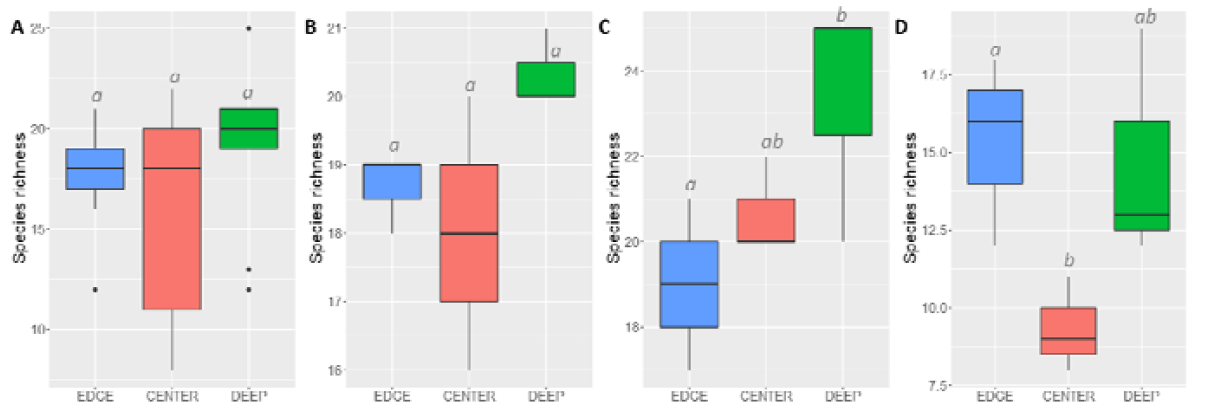
Arthropod species richness (Chao1, q0) in three habitats (forest EDGE, CENTER and DEEP) of Terra-Brava native forest fragment. Boxplots present species richness data for all species A), and for the three biogeographic groups: endemic B), native non-endemic, C) and introduced species D).

**Fig. 5.**
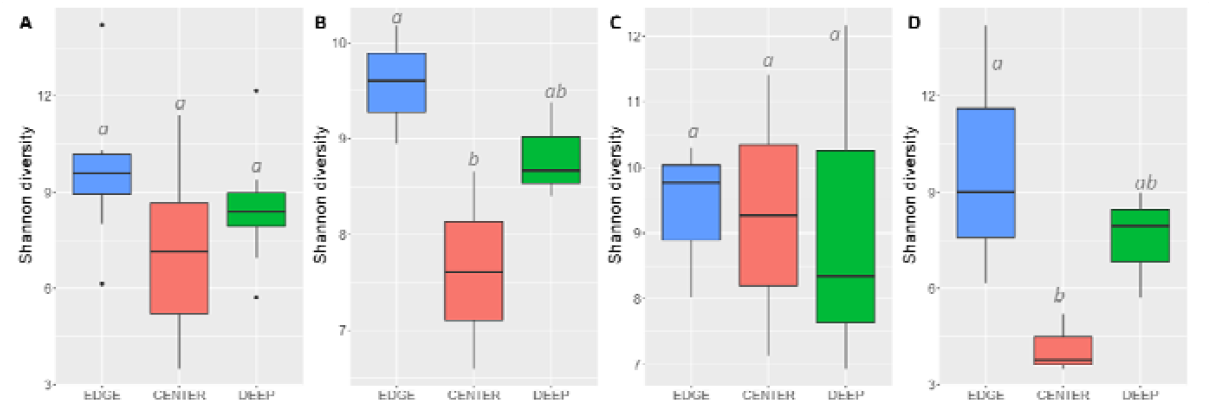
Arthropod species diversity (Shannon diversity, q1) in three habitats (forest EDGE, CENTER and DEEP) of Terra-Brava native forest fragment. Boxplots present Shannon diversity data for all species A), and for the three biogeographic groups: endemic B), native non-endemic, C) and introduced species D).

**Fig. 6.**
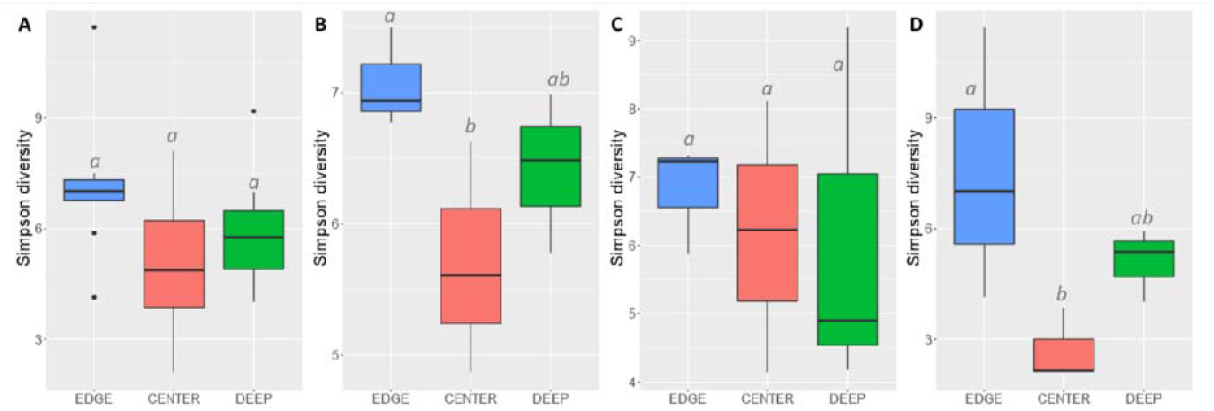
Arthropod species diversity (Simpson diversity, q2) in three habitats (forest EDGE, CENTER and DEEP) of Terra-Brava native forest fragment. Boxplots present Simpson diversity data for all species A), and for the three biogeographic groups: endemic B), native non-endemic, C) and introduced species D).

A higher number of endemic and native species was found in the DEEP areas of the forest compared to outer habitats (Fig. 4B and 4C), but only some differences were significant (Table A.3). Introduced species, however, showed high species richness in EDGE and DEEP habitats being significantly different from the low values recorded in the CENTER habitat (Fig. 4D; Table A.3).

Shannon and Simpson diversity values were similar between all three habitats for native non-endemic species (Fig. 5 and Fig 6; Table A.4, and Table A.5) but both for endemic and introduced species, these two indices had significantly lower values in CENTER than in the DEEP and EDGE habitats (Fig. 5B, 5D, 6B, 6D).

### 3.3. Edge effect on species composition

Models derived from partial dbRDAs (i.e. those with sampling year and sampling month partialled out) were highly significant (p ≤ 0.001) when all species were considered in the dataset as well as when biogeographical categories were analyzed separately.

Similarly, the effect of habitat was also highly significant in all cases (Table A.6). These models explained a lower amount of variance and their AICs varied amongst the four groups (Table 1). The model with all species had the best R^2^ explaining up to 7% of the variance and the model with native non-endemic species explained only 2% of the total variance (Table 1).

**Table 1.**
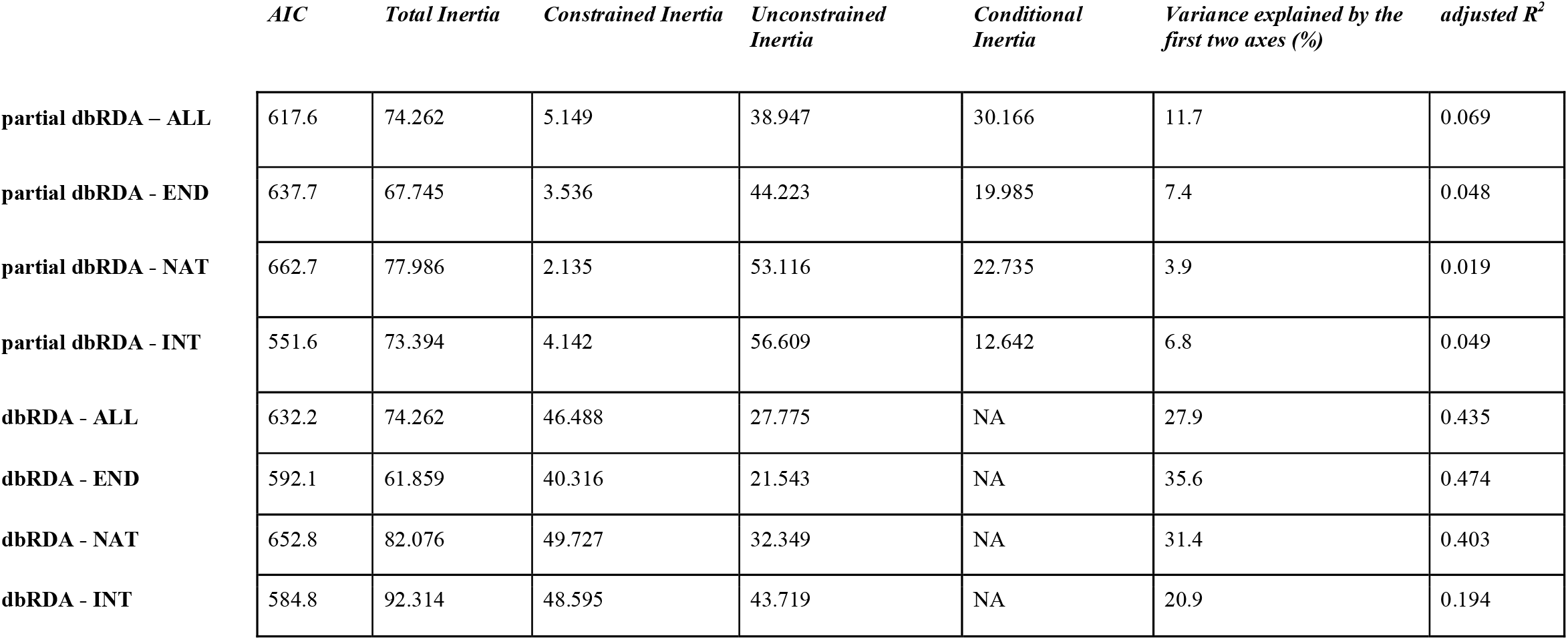
Main characteristics of the dbRDA models with sampling year and sampling month partialled out (partial dbRDA) or included as interacting variables (dbRDA) for ies (ALL), endemic species (END), native non-endemic species (NAT) and introduced species (INT). Species were sampled in the native forest fragment Terra-Brava habitats from border (EDGE) to the core forest (DEEP) and in an intermediate habitat (CENTER).

Arthropod assemblages were well-separated between the three habitats when all species and only endemics were considered (Figure 7A-B), but the similarity of species composition between habitats was high for native non-endemic species and even higher for introduced species (Figure 7C-D).

**Fig. 7.**
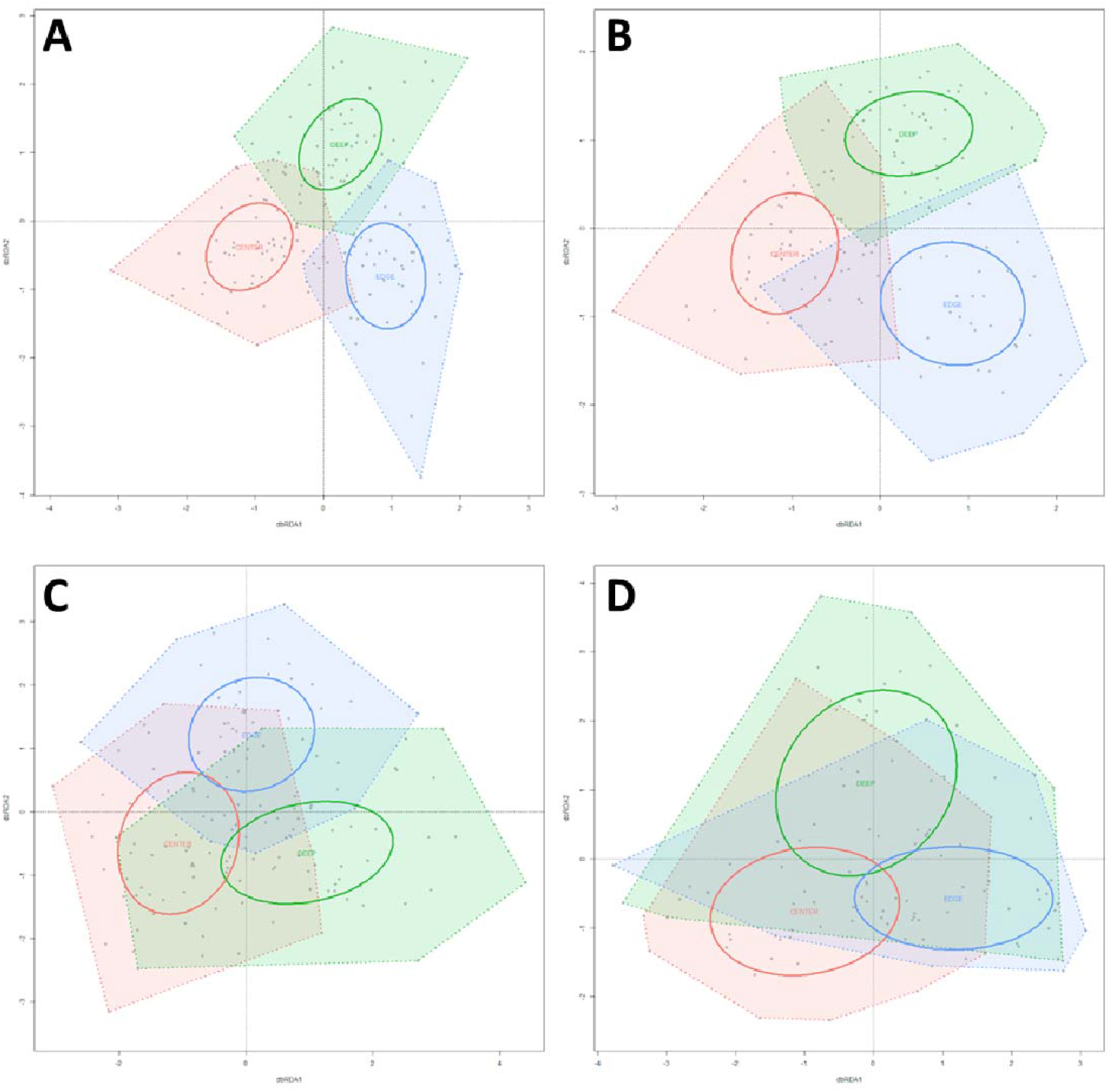
Ordination plots of the partial dbRDA models, in which sampling year and month were partialled out (partial dbRDA), run on all species A), and for the three biogeographic groups: endemic B), native non-endemic, C) and introduced species D). Habitats are color coded: green indicates the deep forest (DEEP), red the intermediate habitat (CENTER), and blue the forest edge (EDGE). Individual sampling sites are indicated with gray dots. Ellipses indicate the standard deviation of the site points belonging to one habitat, and polygons encircle all sites belonging to one habitat.

### 3.4. Edge effect is modulated by species temporal dynamics

#### Abundance

Overall species abundance showed significant variation across the sampling period presenting a unimodal distribution with a peak in summer (August) (Fig.8A). This temporal pattern in arthropod abundance was also recorded for all three biogeographic species groups (Fig.8B-C-D) with slight differences in the exact month of peak abundance. For all species, endemic and native non-endemic species groups, the peak was recorded between July and August in the three habitats (Fig. A.2 panels A-B and C).

**Fig. 8.**
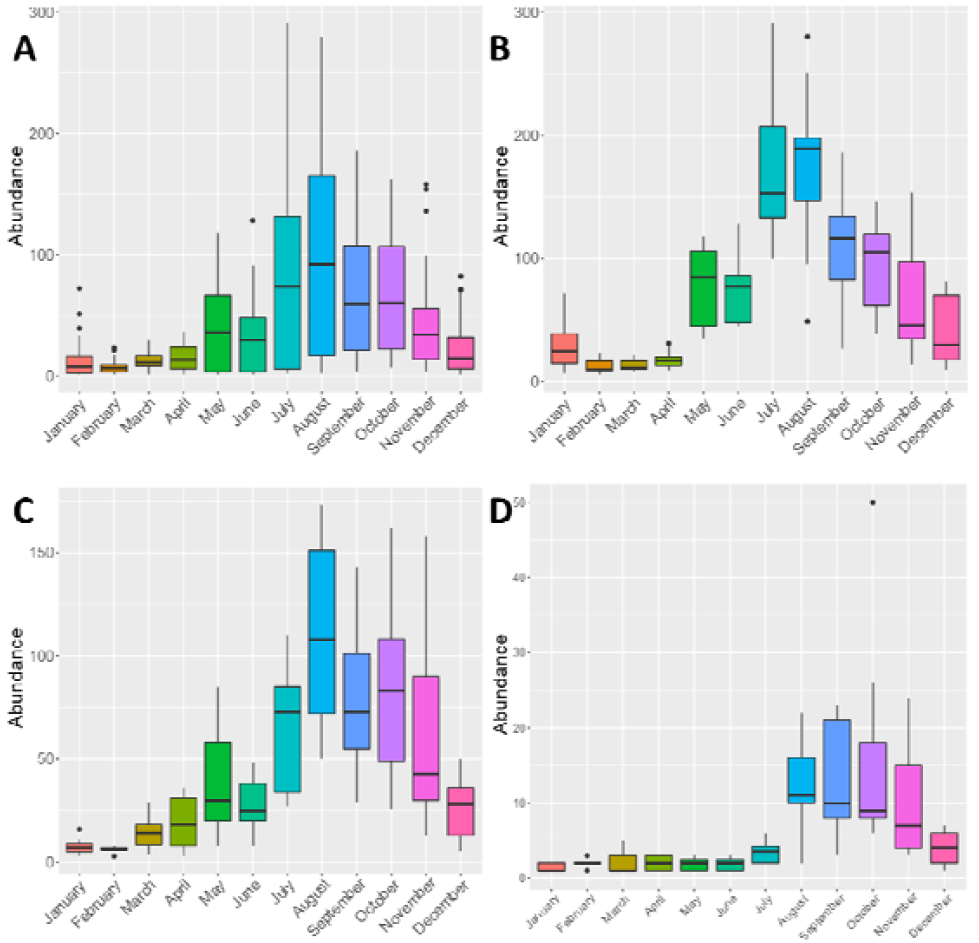
Total arthropod abundance sampled per month in Terra-Brava native forest fragment. Boxplots present abundance data for all species A), and for the three biogeographic groups: endemic B), native non-endemic, C) and introduced species D).

For introduced species, the peak was observed later (in October) in EDGE habitat and between August and October in CENTER and DEEP forest habitats (Fig. A.2 panel D).

The effects of interaction between habitat and month on arthropod abundance was significant during the year (except in autumn) for all species (Table 2) and for endemics (Table A.8), whereas for native non-endemics, significant interactions were observed only in spring (March and April) (Table A.9). For introduced species the effects of the interaction were not significant in any month (Table A.10) which reflects the similarity observed in arthropod abundance between the three habitats (Fig. 3D).

**Table 2.** Summary of the GLMM of total arthropods abundance for all species. Species were sampled in the native forest fragment Terra-Brava in three habitats from border (EDGE) to the core forest (DEEP) and in an intermediate habitat (CENTER). Variance explained by fixed effects: Marginal R^2^ = 0.91. Variance explained by the entire model: Conditional R^2^ = 0.93.

|  | *Estimate* | *Std. Error* | *z value* | *P* |
| --- | --- | --- | --- | --- |
| **Habitat** |  |  |  |  |
| Intercept | 1.864 | 0.155 | 11.993 | <0.0001 |
| Center | 0.559 | 0.196 | 2.850 | 0.004 |
| Deep | 1.184 | 0.184 | 6.423 | <0.0001 |
| **Month** |  |  |  |  |
| February | 0.202 | 0.188 | 1.076 | 0.282 |
| March | 0.633 | 0.177 | 3.586 | <0.001 |
| April | 0.981 | 0.164 | 5.973 | <0.0001 |
| May | 1.693 | 0.152 | 11.127 | <0.0001 |
| June | 1.502 | 0.153 | 9.814 | <0.0001 |
| July | 1.699 | 0.146 | 11.617 | <0.0001 |
| August | 1.942 | 0.145 | 13.364 | <0.0001 |
| September | 1.597 | 0.148 | 10.802 | <0.0001 |
| October | 1.230 | 0.149 | 8.268 | <0.0001 |
| November | 0.972 | 0.153 | 6.339 | <0.0001 |
| December | 0.221 | 0.171 | 1.293 | 0.196 |
| **Habitat × Month** |  |  |  |  |
| Center*February | -0.834 | 0.256 | -3.263 | <0.01 |
| Center*March | -0.920 | 0.243 | -3.787 | <0.001 |
| Center*April | -1.279 | 0.232 | -5.506 | <0.0001 |
| Center*May | -0.704 | 0.193 | -3.658 | <0.001 |
| Center*June | -0.321 | 0.192 | -1.666 | 0.096 |
| Center*July | -0.410 | 0.181 | -2.269 | 0.023 |
| Center*August | -0.573 | 0.179 | -3.196 | <0.01 |
| Center*September | -0.753 | 0.183 | -4.105 | <0.0001 |
| Center*October | -0.200 | 0.183 | -1.091 | 0.275 |
| Center*November | -0.911 | 0.194 | -4.685 | <0.0001 |
| Center*December | -0.443 | 0.214 | -2.072 | 0.038 |
| Deep*February | -1.121 | 0.235 | -4.769 | <0.0001 |
| Deep*March | -1.021 | 0.215 | -4.745 | <0.0001 |
| Deep*April | -1.069 | 0.197 | -5.440 | <0.0001 |
| Deep*May | -0.630 | 0.174 | -3.625 | <0.001 |
| Deep*June | -0.719 | 0.176 | -4.075 | <0.0001 |
| Deep*July | -0.675 | 0.166 | -4.059 | <0.0001 |
| Deep*August | -0.774 | 0.165 | -4.693 | <0.0001 |
| Deep*September | -0.765 | 0.168 | -4.551 | <0.0001 |
| Deep*October | -0.472 | 0.169 | -2.788 | 0.005 |
| Deep*November | -0.214 | 0.173 | -1.235 | 0.217 |
| Deep*December | -0.186 | 0.192 | -0.968 | 0.333 |

#### Species richness

Arthropod species richness also varied across the sampling period with a peak during summer months: July, August and September (Fig. 9A). For endemic species, species richness increased progressively from April and peaked in July, smoothly decreasing thenceforth (Fig. 9B). A similar pattern was observed for native non-endemic species (Fig. 9C) while the introduced species distribution showed one peak in October (Fig. 9D). When considering the distinct habitats, for all species, for endemic and for native non-endemic species groups, the peak was recorded between July and August (Fig. A.3 panels A-B and C) and for introduced species the peaks were early Autumn (Fig. A.3 panel D).

**Fig. 9.**
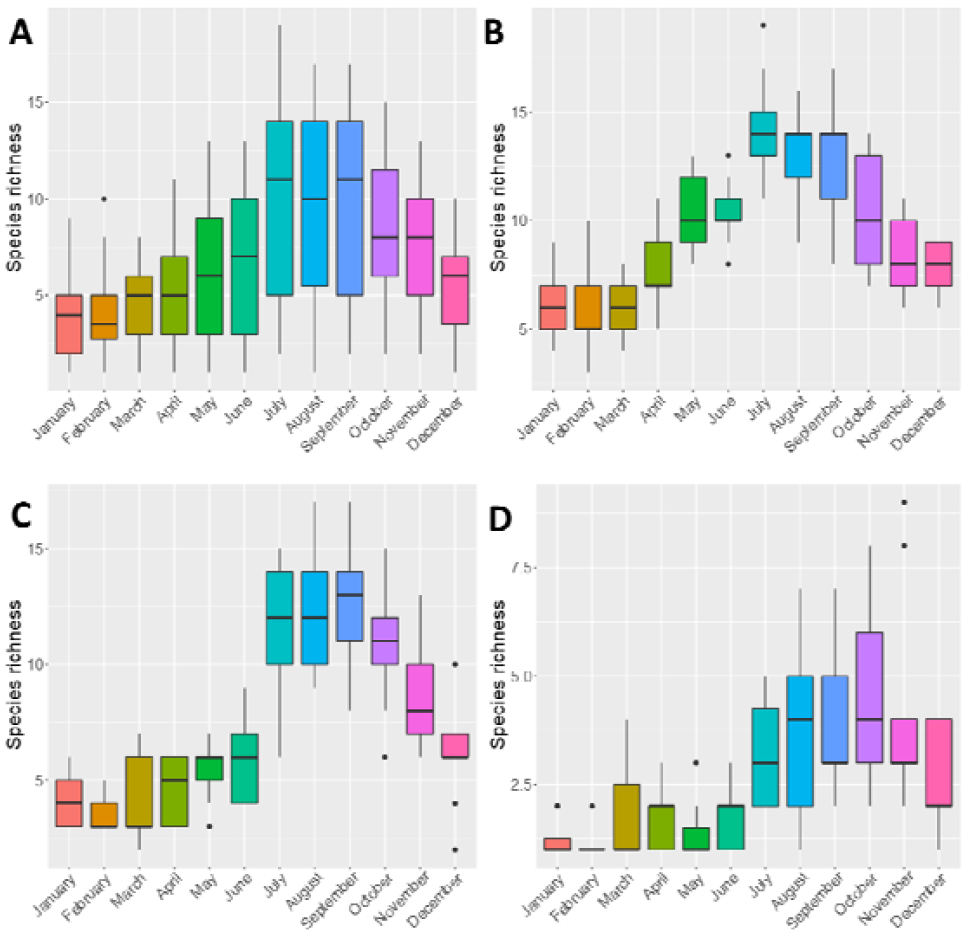
Total arthropod species richness (chao1, q0) sampled per month in Terra-Brava native forest fragment. Boxplots present species richness data for all species A), and for the three biogeographic groups: endemic B), native non-endemic C) and introduced species D).

We found a significant effect of the sampling period (month) in late summer for endemic and native non-endemic species (July to Oct, Table A.11 and A.12; Fig. A.3 panels A-B and C), and during early autumn (Oct, Nov, Table A.13) for introduced species. However, models did not show significant effects of habitat nor interaction of habitat and month (Table 3, A.11-13; Fig. A.3 panel D).

**Table 3.** Summary of the GLMM of total arthropods richness (Chao1, q0) for all species. Species were sampled in the native forest fragment Terra-Brava in three habitats from border (EDGE) to the core forest (DEEP) and in an intermediate habitat (CENTER). Variance explained by fixed effects: Marginal R2 = 0.44. Variance explained by the entire model: Conditional R2 = 0.45.

|  | *Estimate* | *Std. Error* | *z value* | *P* |
| --- | --- | --- | --- | --- |
| **Habitat** |  |  |  |  |
| Intercept | 1.292 | 0.189 | 6.852 | <0.0001 |
| Center | 0.006 | 0.259 | 0.022 | 0.982 |
| Deep | 0.199 | 0.248 | 0.800 | 0.424 |
| **Month** |  |  |  |  |
| February | 0.028 | 0.261 | 0.109 | 0.913 |
| March | 0.281 | 0.253 | 1.110 | 0.267 |
| April | 0.417 | 0.239 | 1.743 | 0.081 |
| May | 0.534 | 0.234 | 2.284 | 0.022 |
| June | 0.381 | 0.235 | 1.618 | 0.106 |
| July | 0.950 | 0.219 | 4.345 | <0.0001 |
| August | 0.916 | 0.216 | 4.239 | <0.0001 |
| September | 0.916 | 0.216 | 4.239 | <0.0001 |
| October | 0.840 | 0.218 | 3.848 | <0.001 |
| November | 0.604 | 0.226 | 2.669 | 0.008 |
| December | 0.316 | 0.238 | 1.327 | 0.185 |
| **Habitat × Month** |  |  |  |  |
| Center*February | 0.029 | 0.361 | 0.079 | 0.937 |
| Center*March | -0.130 | 0.365 | -0.356 | 0.722 |
| Center*April | -0.220 | 0.346 | -0.636 | 0.525 |
| Center*May | -0.015 | 0.325 | -0.047 | 0.962 |
| Center*June | 0.284 | 0.325 | 0.874 | 0.382 |
| Center*July | -0.090 | 0.302 | -0.298 | 0.765 |
| Center*August | -0.122 | 0.301 | -0.406 | 0.685 |
| Center*September | -0.043 | 0.299 | -0.144 | 0.885 |
| Center*October | -0.226 | 0.307 | -0.735 | 0.462 |
| Center*November | -0.130 | 0.317 | -0.410 | 0.682 |
| Center*December | -0.099 | 0.334 | -0.297 | 0.767 |
| Deep*February | -0.169 | 0.354 | -0.478 | 0.633 |
| Deep*March | -0.198 | 0.344 | -0.575 | 0.565 |
| Deep*April | -0.185 | 0.323 | -0.572 | 0.567 |
| Deep*May | -0.215 | 0.313 | -0.688 | 0.491 |
| Deep*June | 0.041 | 0.311 | 0.133 | 0.894 |
| Deep*July | 0.015 | 0.287 | 0.052 | 0.959 |
| Deep*August | 0.077 | 0.285 | 0.271 | 0.786 |
| Deep*September | 0.010 | 0.286 | 0.035 | 0.972 |
| Deep*October | 0.035 | 0.288 | 0.123 | 0.902 |
| Deep*November | 0.089 | 0.298 | 0.300 | 0.764 |
| Deep*December | 0.106 | 0.313 | 0.338 | 0.735 |

#### Species diversity

Arthropod species diversity (measured by both Shannon and Simpson diversity indices) also varied across the sampling period. The data set for all species showed moderate fluctuation in diversity throughout the year while native non-endemic species had a considerable increase in diversity in July followed by a steady decline in October (Fig. 10AC and Fig 11AC). Diversity showed a bimodal distribution for endemic species, peaking in April and in September (Fig. 10B and Fig. 11B) while for introduced species, diversity distribution showed two plateaus, a lower from January to June and an upper from the rest of the year (Fig. 10A and Fig 11A).

**Fig. 10.**
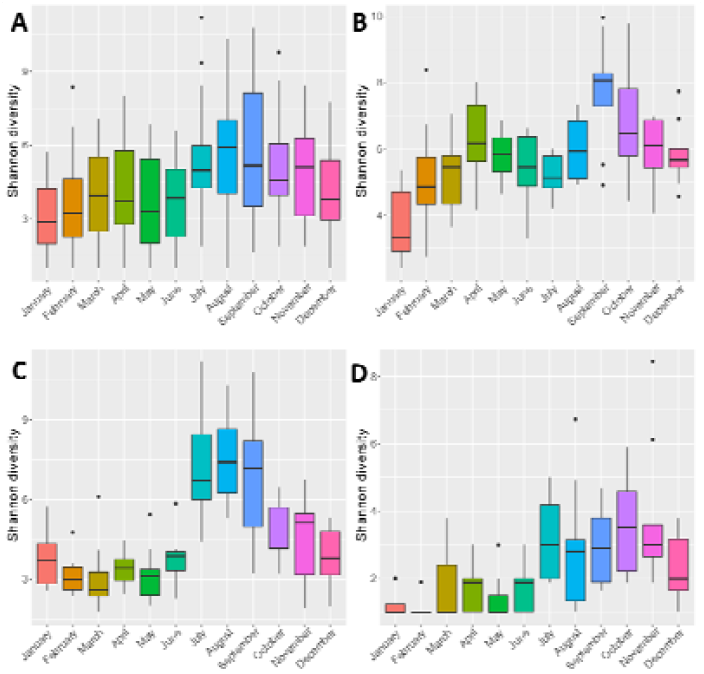
Total arthropod species diversity (Shannon diversity, q1) sampled per month in Terra-Brava native forest fragment. Boxplots present Shannon diversity data for all species A), and for the three biogeographic groups: endemic B), native non-endemic C) and introduced species D).

**Fig. 11.**
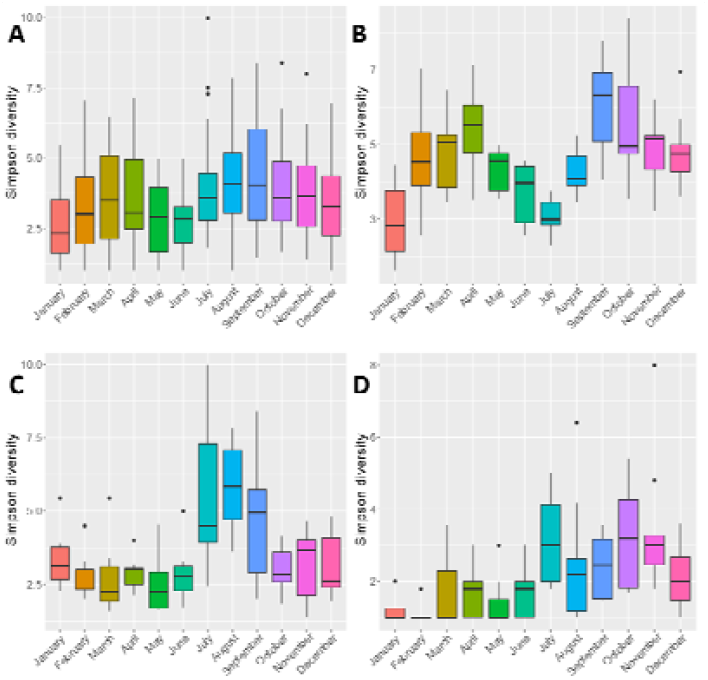
Total arthropod species diversity (Simpson diversity, q2) sampled per month in Terra-Brava native forest fragment. Boxplots present Simpson diversity data for all species A), and for the three biogeographic groups: endemic B), native non-endemic C) and introduced species D).

Models showed significant effects of both habitat and sampling month on species diversity (Tables 4 and 5). Diversity peaks were observed between June and September for all species, endemic and native non-endemic species in the three habitats (Fig. A.4 and A.5 panels A, B and C) whereas for introduced species peaks were observed in October in edge habitat, between August and November in center habitat and between July and November in deep habitat (Fig. A.4 and A.5 panel D). The interaction of the two factors was not significant neither for all species, nor for the three biogeographic groups (Tables A.14-19).

**Table 4.**
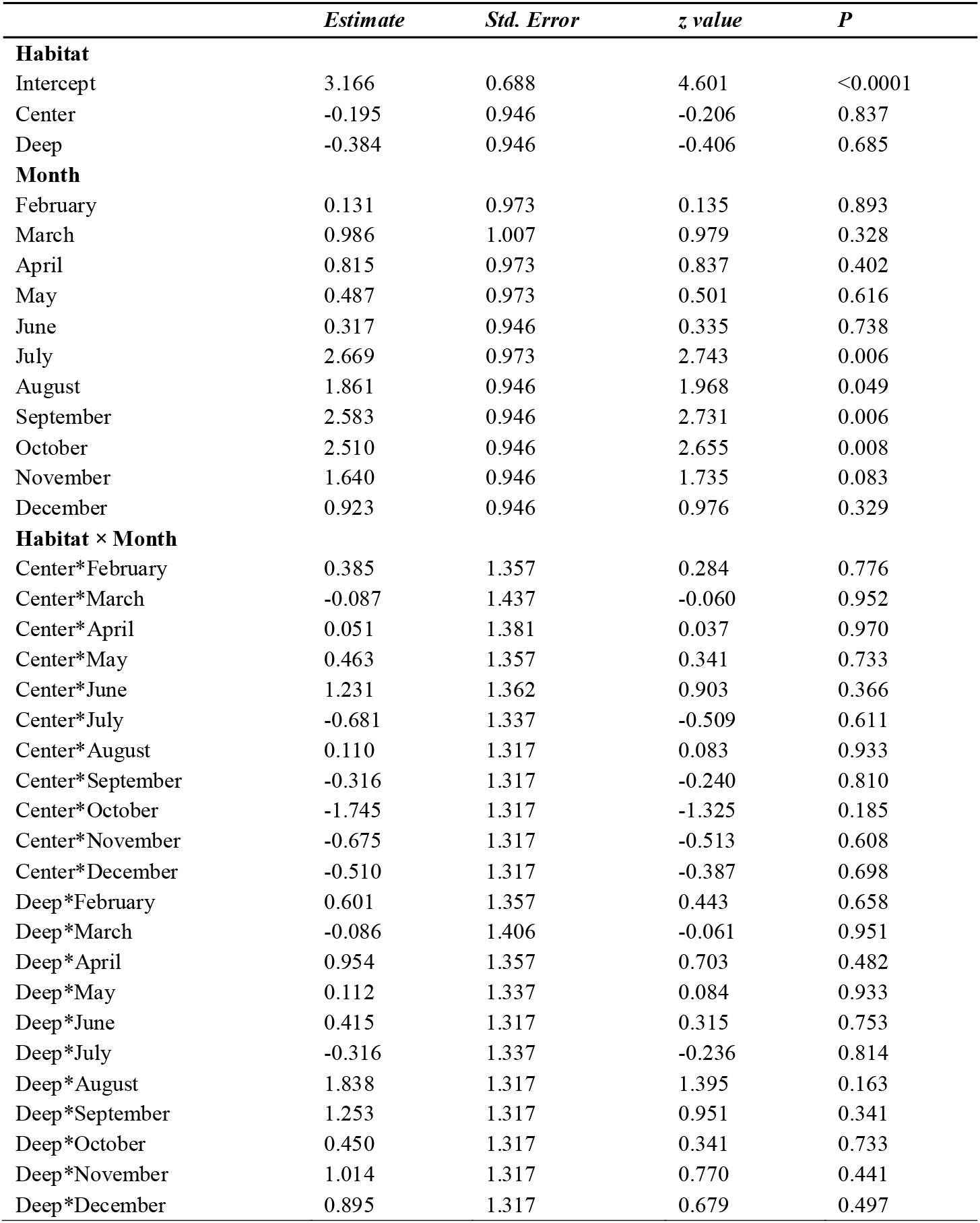
Summary of the GLMM of total arthropods species diversity (Shannon diversity, q1) for all species. Species were sampled in the native forest fragment Terra-Brava in three habitats from border (EDGE) to the core forest (DEEP) and in an intermediate habitat (CENTER). Variance explained by fixed effects: Marginal R^2^= 0.21. Variance explained by the entire model: Conditional R^2^= 0.21.

**Table 5.** Summary of the GLMM of total arthropods species diversity (Simpson diversity, q2) for all species. Species were sampled in the native forest fragment Terra-Brava in three habitats from border (EDGE) to the core forest (DEEP) and in an intermediate habitat (CENTER). Variance explained by fixed effects: Marginal R^2^ = 0.11. Variance explained by the entire model: Conditional R^2^ = 0.11.

|  | *Estimate* | *Std. Error* | *z value* | *P* |
| --- | --- | --- | --- | --- |
| **Habitat** |  |  |  |  |
| Intercept | 1.036 | 0.211 | 4.917 | <0.0001 |
| Center | -0.051 | 0.293 | -0.175 | 0.861 |
| Deep | -0.238 | 0.307 | -0.775 | 0.438 |
| **Month** |  |  |  |  |
| February | 0.049 | 0.294 | 0.168 | 0.867 |
| March | 0.288 | 0.287 | 1.002 | 0.316 |
| April | 0.147 | 0.288 | 0.511 | 0.609 |
| May | 0.004 | 0.298 | 0.013 | 0.990 |
| June | -0.009 | 0.290 | -0.030 | 0.976 |
| July | 0.489 | 0.268 | 1.827 | 0.068 |
| August | 0.260 | 0.273 | 0.951 | 0.342 |
| September | 0.452 | 0.264 | 1.716 | 0.086 |
| October | 0.501 | 0.261 | 1.919 | 0.055 |
| November | 0.336 | 0.269 | 1.248 | 0.212 |
| December | 0.226 | 0.275 | 0.822 | 0.411 |
| **Habitat × Month** |  |  |  |  |
| Center*February | 0.114 | 0.410 | 0.278 | 0.781 |
| Center*March | -0.006 | 0.413 | -0.014 | 0.989 |
| Center*April | 0.092 | 0.408 | 0.227 | 0.821 |
| Center*May | 0.133 | 0.413 | 0.321 | 0.748 |
| Center*June | 0.237 | 0.410 | 0.578 | 0.564 |
| Center*July | -0.139 | 0.377 | -0.369 | 0.712 |
| Center*August | 0.135 | 0.380 | 0.356 | 0.722 |
| Center*September | -0.074 | 0.373 | -0.199 | 0.842 |
| Center*October | -0.425 | 0.385 | -1.104 | 0.270 |
| Center*November | -0.182 | 0.387 | -0.471 | 0.638 |
| Center*December | -0.169 | 0.396 | -0.426 | 0.670 |
| Deep*February | 0.336 | 0.418 | 0.803 | 0.422 |
| Deep*March | 0.055 | 0.422 | 0.130 | 0.897 |
| Deep*April | 0.449 | 0.405 | 1.109 | 0.267 |
| Deep*May | 0.167 | 0.425 | 0.393 | 0.694 |
| Deep*June | 0.114 | 0.423 | 0.268 | 0.788 |
| Deep*July | -0.013 | 0.391 | -0.033 | 0.973 |
| Deep*August | 0.563 | 0.383 | 1.470 | 0.142 |
| Deep*September | 0.362 | 0.376 | 0.962 | 0.336 |
| Deep*October | 0.137 | 0.380 | 0.359 | 0.720 |
| Deep*November | 0.311 | 0.386 | 0.806 | 0.420 |
| Deep*December | 0.277 | 0.395 | 0.702 | 0.483 |

#### Species composition

Standard dbRDA models, with sampling year and sampling month included as interacting variables, were also all significant, and showed mostly significant (0.01 ≤ p ≤ 0.05) or highly significant (p ≤ 0.01) effects of habitat, sampling month, and sampling year, as well as their interactions except between the full sampling date (i.e. year and month) and habitat (Table A.7). The combined effect of sampling year and month was not significant either for the endemic or the introduced groups. For this latter group the effect of sampling year and the combined effect of sampling year and habitat were not significant either (Table A.20). Models were robust explaining up to 36% of the variance in endemic species model and up to 21% for introduced species model. Models R^2^ were also relatively high (47% in endemic species model and 20% in introduced species model) (Table 1).

The standard dbRDAs showed a larger overlap between habitats and also indicated a grouping between the EDGE and DEEP forest habitat, and a separation of the CENTER habitat from these two. This pattern was visible when all species were included in the analysis and for the endemics (Figure 12A-B) but seemingly disappeared for the native non-endemic and introduced species (Figure 12C-D).

**Figure 12.**
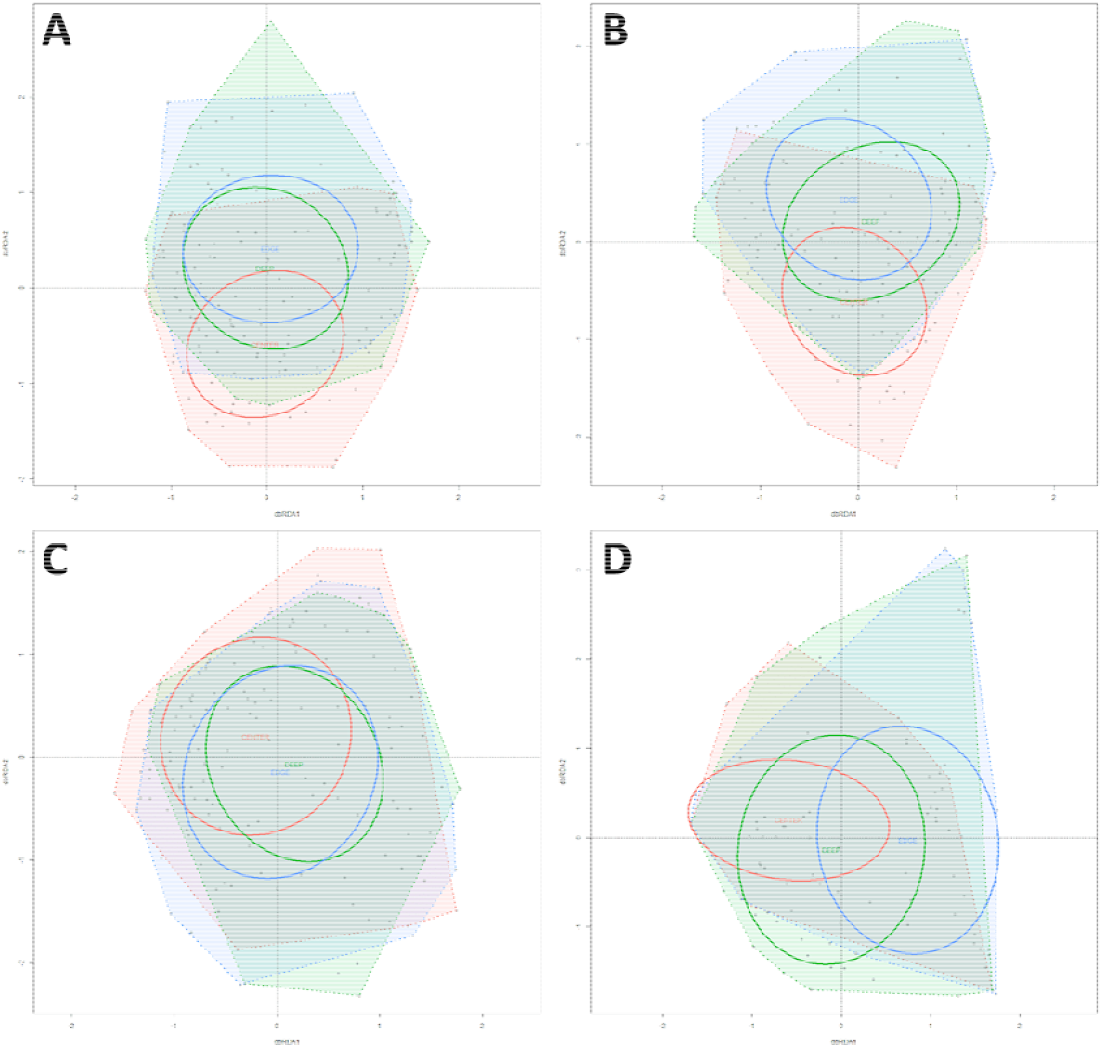
Ordination plots of the standard dbRDA models, in which sampling year and month were included as interacting factors, run on all species A), and for the three biogeographic groups: endemic B), native non-endemic, C) and introduced species D). Habitats are color coded: green indicates the deep forest (DEEP), red the intermediate habitat (CENTER), and blue the forest edge (EDGE). Individual sampling sites are indicated with grey diamonds. Ellipses indicate the standard deviation of the site points belonging to one habitat, and polygons encircle all sites belonging to one habitat.

The NMDS ordination of the samples separated by sampling months and habitats indicated a seasonal pattern, with deep forest and forest edge habitats following a similar trajectory, whereas samples from the intermediate habitat separated well from the other two in most of the year, only with winter samples (from December-February) showing some similarity (Fig. 13). Euclidean distances of the assemblages also showed a similar temporal pattern both within and between samples (i.e. variation within a month, and difference between months, respectively), with greater mean distances both within- and between-habitat samples early in the year, a homogenization from spring (May) and, with some variation, gradually increasing distances again from August-September (Fig. A.6). The range of the within-sample distances (i.e. distances from samples taken from the same habitat in the same month) tended to be lower in the late spring - early summer (Fig. A.6).

**Figure 13.**
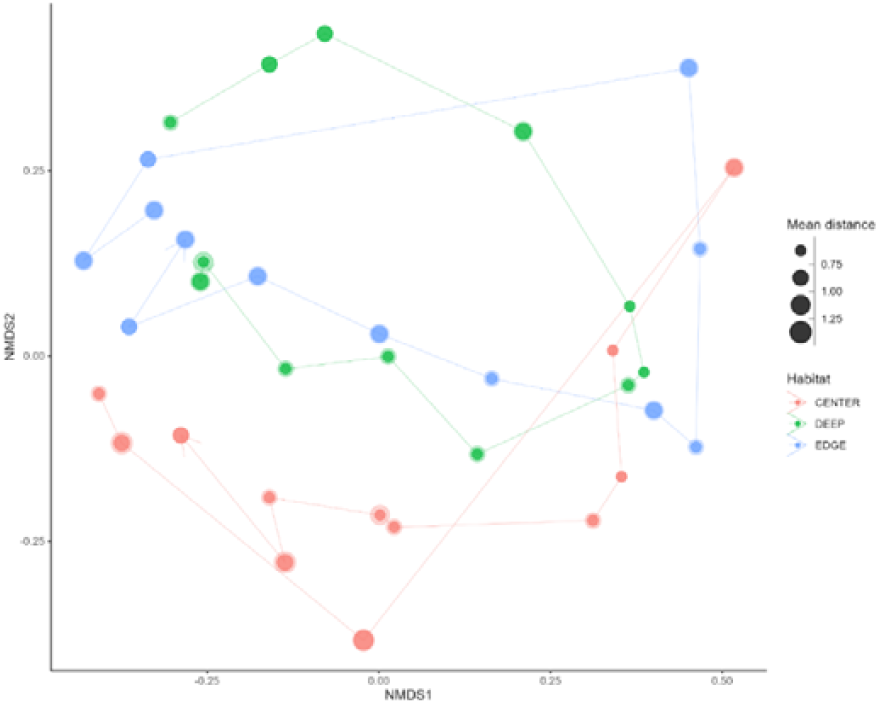
Multidimensional scaling of monthly samples in the three habitats showing the seasonal trajectory of between-month similarities. Habitats are color coded: green indicates the deep forest (DEEP), red the intermediate habitat (CENTER), and blue the forest edge (EDGE). The size of the points is proportional to the within-month mean distance between samples, and the shading around points indicates the standard deviation.

## 4. Discussion

In our goal to test the edge effect in native Azorean forests, we investigated how arthropod communities differed between edge, intermediate, and deep forest habitats, with a particular interest in assessing the different responses of endemic, native non-endemic and introduced species. We found evidence for changes in biodiversity due to edge effects, but these responses varied temporally and according to species’ biogeographic categories.

Previous studies on the arthropod fauna of Azorean native forest canopies showed that endemic and native non-endemic species dominated the assemblages while introduced species occur in very low overall abundance (Ribeiro et al., 2005; Gaspar et al., 2011; Rego et al., 2019). For example, Gaspar et al. (2011) reported that most individuals sampled in Azorean native forest fragments were endemic species (49%) but, endemic (108 spp) and native non-endemic species richness (149spp) were significantly lower than that of introduced species (195 spp). Similarly, Rego et al. (2019), also found that indigenous species (endemic and native non-endemic species) were more abundant than introduced species, counting for about 68% of the total abundance. Yet, we found that species numbers were similarly distributed within the three biogeographic categories, suggesting that there is a trend of a continuous rain of introduced species arriving in the canopies of Azorean native forests but still not establishing abundant populations. This source-sink dynamics is promoting an increase on the diversity of exotic arthropods in the Azorean native forest through time and is a matter of conservation concern (see Borges et al., 2020).

### 4.1. Edge effects determine species abundances

In agreement with our expectations, arthropod abundances were higher in forest interior than at edges or in intermediates habitats and these results suggest that arthropod assemblages were sensitive to distance to the edge. Since most collected specimens belonged to indigenous species, whose number may drive overall abundance patterns, it is plausible to assume that greater insect abundances in the forest interior were observed because of the abundance of native plants to which indigenous insects are tightly linked (Rego et al., 2019). In fact, our results also showed that while the abundance of indigenous species increased from the edge to the core forest, introduced species were similarly abundant in all three habitats. Introduced species are less likely to be associated with native trees and are more likely to be generalist species, which can explain why their abundance is unimpacted by either the edge effect or habitat quality in general. Another explanation is the fact that some of the exotic species are spiders (like *Ero furcata* and *Tenuiphantes tenuis)* that perform ballooning, enabling them to enter the canopies of forest at all places and consequently diminishing the influence of edge effect.

Unlike our multitaxon approach, most studies (but see Jokimäki et al., 1998) investigated the edge effect on a particular taxonomic group. Some of these taxonomically more restricted approaches showed similar patterns to our results. For instance, a decrease in dung beetle abundance was observed near to edges in a forest – sun-grown coffee ecotone (Villada-Bedoya et al., 2017) and a similar pattern was observed in Afromontane rainforests (Barnes et al., 2014). Yet, as reviewed by Stone et al. (2018), most studies observed no edge effect on arthropod abundances, and a few even reported an increase in species abundance near to edges (Lacasella et al., 2015).

Our results indicate that arthropods were as abundant at edges as in the intermediate habitat. This suggests that the edge effect on abundance in our study is extended over a distance of 100 m to forest interior for indigenous species while the few introduced individuals we observed (4% of total abundance) seem to be well supported at edges as well as in forest interior and therefore not sensitive to edge effect. As most of these exotics introduced species are likely to be ‘tourists’, they presumably are enrolled in a source-sink dynamic and consequently most of their populations are unlikely to be in equilibrium in the native forest (Borges et al., 2005).

### 4.2. Edge effects do not impact species richness or diversity

We expected to find an increase of indigenous species from the edge toward forest interior and, in turn, more introduced species at edges compared to forest interior. Considering all species together, our results did not support these expectations; rather, we found no differences in richness and diversity indices between the three habitats. This suggests that even if species realized their niches at the interior of the forest (significant higher species abundance at the forest interior), they can move from one habitat to another without a barrier. This was true for rare (Hill number q = 0) as well as for common (q = 1) and dominant (q = 2) species. This result is not surprising as most of the collected individuals (89%) and species (84%) sampled were species with high dispersal ability.

Contrasting results were however observed when we considered biogeographical categories. In this case, we surprisingly did not observe an edge effect but an “intermediate effect” within introduced species assemblages. Introduced species richness was lower at intermediate habitat but similar between edges and forest interior habitats. Moreover, this pattern was observed equally for rare, common and dominant species. Patterns were different between endemic and native non-endemic species: we found that common and dominant endemic species showed the “intermediate effect” that we observed for introduced species, yet in native non-endemic species, such effect was observed only in rare species but neither for common nor for dominant species.

Contrasting to our results, clear relationships between species richness and distance from forest edges were found in beetle assemblages. For instance Magura (2002) found higher carabid species richness near to the forest edge in a forest-grassland ecotone whereas Stone et al. (2018) reported the opposite pattern, a decline of beetles species richness toward a sclerophyllous forest interiors adjacent to suburban areas. However, our study encompasses more diverse arthropod assemblage (15 orders) which may be one of the reasons for the differences from the aforementioned studies. Moreover, our study system was in a humid native forest adjacent semi-natural pasture to an exotic forest (dominated by *Cryptomoria japonica, Pittosporum undulatum, Eucalyptus* spp trees species) where vegetation cover between the edge and forest interior was less obvious than in the aforementioned studies (Magura, 2002; Stone et al., 2018).

### 4.3. Edge effect on species composition

The composition of species seems to be impacted by forest edge, with the habitat type significantly determining the species composition not only of the community at large but also of species assemblages of different biogeography categories, particularly those of endemic species. Arthropod species assemblages of the three habitats were completely separated when all species and endemics were considered. This result indicates that the composition of endemic species was specific to habitat and probably being constrained by local habitat characteristics. For example, the endemic rove beetle *A. dryochares* seems to be restricted to the inner areas of the forest, where hyper-humid and more pristine conditions prevail. In contrast, the endemic cicadellid *Eupteryx azorica* was associated with forest edges habitats, as was the endemic spider *Canariphantes acoreensis* which builds its webs on less humid and more sunlit environments.

Although the effect of habitat was also significant when native non-endemic or introduced species were considered, the separation of species assemblages was less clear, with a slight overlap in native non-endemic species and a high overlap in introduced species. These structural and compositional differences between habitats drive the segregation of several species along the gradient from the edge to the inner forest. This finding suggests that there is an increase of habitat specificity from endemic to native non-endemic and introduced species community.

### 4.4 Edge effect is mediated by species temporal dynamic

Since most species follow seasonal patterns in their activity (Borges et al., 2017), we predicted that arthropod communities will vary in time and their responses to edge effects will be mediated by the species’ temporal dynamics. Our results partially validated this hypothesis. Univariate models showed a significant interactive effect of habitat, and sampling period on species abundance, confirming that edge effect determines species abundance and that the effect depends on the season considered. Nonetheless, we found that the biogeographical categories also play an important role. Although endemic species replicated the trend anticipated from the arthropod activity patterns on the island (Gaspar et al., 2008), native non-endemic and introduced species abundances seem to be less sensitive to edge effect at any time in the year. Conversely, we found no temporal effect on species richness or diversity. The dbRDA ordination plots showed high overlap in species composition of the three habitats indicating that the composition of species in the three habitats is similar for all species as well as for the three biogeographic categories.

### 4.5. Implication for management

Our study exposed effects forest edges have on arthropod communities collected inside and at the border of a pristine island native forest. The effects were mediated by species biogeographic categories and temporal dynamics. We found that the abundance of endemic species was strongly constrained by edges and that, even if some species are located nearer edges, most of them are more abundant at the core forest interior. Our results also revealed that the edge can have an effect on endemic species for up to 100 m inside the forest. This distance is notable, taking into consideration that many Azorean native forest fragments have a small area, comparable to this distance, which may further contribute to the decline of the already threatened endemic arthropod species. Indeed, many Azorean endemic arthropods are considered as being in extinction debt (Triantis et al., 2010), and some species were recently classified as Critically Endangered or Endangered by the IUCN (see http://www.maiisg.com/). Our findings reiterate how detrimental human impacts, such as increasing fragmentation and edge effects, can be to these endangered organisms.

In contrast, introduced species do not seem to be impacted. Even though they were found in a low relative abundance, introduced species were equally spread and equally diverse at the edges as in forest interior. Here, thus, we highlight how deleterious activities that lead to an increase in edge habitats can be by strongly constraining endemic species but not limiting introduced ones. Indeed, our study demonstrates that edges might favor some species but those species were mostly exotic species, which include several invasive species (e.g. the spider *Dysdera crocata*), and that in contrast endemics were secluded in the forest interior. We can therefore predict that the more the border (the more fragmentation) the less suitable habitat endemic species will have. Although the Azorean authorities are committed to protect the last remaining native forest fragments (mostly of smaller size and vulnerable to edge effects) and restoration initiatives are underway in several islands to increase the area and connectivity of native forest fragments (ongoing Projects LIFE-BEETLES; LIFE-SNAILS and LIFE-IP), the situation may not be as good in other islands.

In the SLOSS debate (Tjørve, 2010) opposing single large reserve strategy to several small reserves strategy, some studies claim that the several small areas strategy was the best to preserve as many species as possible. However, this strategy by definition involves fragmentation and therefore it is not suitable to conserve native and endemic forest species. In order to prevent the impoverishment of native arthropod assemblages, stakeholders should rather focus on minimizing these impacts and increase forest patch size and connectivity.

## Funding

N.T. and M.T.F. are supported by Secretaria Regional do Ambiente e Alterações Climáticas—LIFE-BETTLES (LIFE18 NAT_PT_000864) (2020). M.B. was supported by FCT - DL57/2016/CP1375/CT0001. Field work funded by Portuguese FCT-NETBIOME –ISLANDBIODIV grant 0003/2011 (between 2012 and 2015). The manuscript was also partly financed by FCT–Fundação para aCiência e a Tecnologia, within the project UID/BIA/00329/2013–2020. The Natural Park of Terceira (Azores) provided the necessary authorization for sampling.

## Acknowledgements

We thank the many people who have assisted us with fieldwork, collecting specimens. A large number of students financed by the EU Programs ERASMUS and EURODYSSÉE sorted the samples prior to species assignment by one of us (P.A.V.B.), and we are grateful to all of them: Adal Humberto Díaz Raya, David Rodilla Rivas, Daniel Ehrhart, Juan Ignacio Pitarch Peréz, Juan Manuel Taboada Alvarez, Helena Marugán Páramo, Laura Cáceres Sabater, Laura Gallardo, Marija Tomašic, Óscar García Contreras, Percy de Laminne de Bex, Ruben Murillo Garcia, Rui Carvalho, Sergio Fernandez, Sophie Wallon, and William Razey.

## Data Availability Statement

The data presented in this study are available in Borges et al. (2022) https://doi.org/10.3897/BDJ.10.e85971. Supplementary data are available with the text.

## Author Contributions

Conceptualization, N.T. and P.A.V.B.; data curation, M.B.,; A.R.-P., R.N., and P.A.V.B.; formal analysis, N.T. and G.P.; funding acquisition, M.T. F and P.A.V.B.; investigation, N.T., G.P., M.B., M.T.F., and P.A.V.B.; methodology, N.T., G.P., and P.A.V.B.; project administration, M.T. F and P.A.V.B.; software, N.T. and G.P.; supervision, P.A.V.B.; validation, N.T., G.P., M.B., and P.A.V.B.; writing— original draft, N.T. and G.P.; writing—review and editing, N.T., G.P., M.B., A.R.-P., R.N., M.T.F., and P.A.V.B. All authors have read and agreed to the published version of the manuscript.

## Notes

### Competing Interest Statement

The authors have declared no competing interest.

