## Supplementary data for "Edge effects constraint endemic but not introduced arthropod species in a pristine forest on Terceira (Azores, Portugal)"

^1^cE3c- Centre for Ecology, Evolution and Environmental Changes, Azorean Biodiversity Group, CHANGE – Global Change and Sustainability Institute, Faculty of Agricultural Sciences and Environment, University of the Azores, Rua Capitão João d´Ávila, Pico da Urze, 9700-042, Angra do Heroísmo, Portugal. ^2^Regional Secretariat of Environment and Climate Change, Project LIFE BEETLES (LIFE 18NAT/PT/000864), Rua do Galo n118, 9700-040 Angra do Heroísmo, Açores, Portugal. ^3^IUCN SSC Mid-Atlantic Islands Invertebrates Specialist Group, Angra do Heroísmo, Azores, Portugal

Authors emails:

G.P.:

M.B.:

A.R-P.:

R.N.:

M.T.F.:

P.A.V.B.:

### Supplementary Material

#### Figures

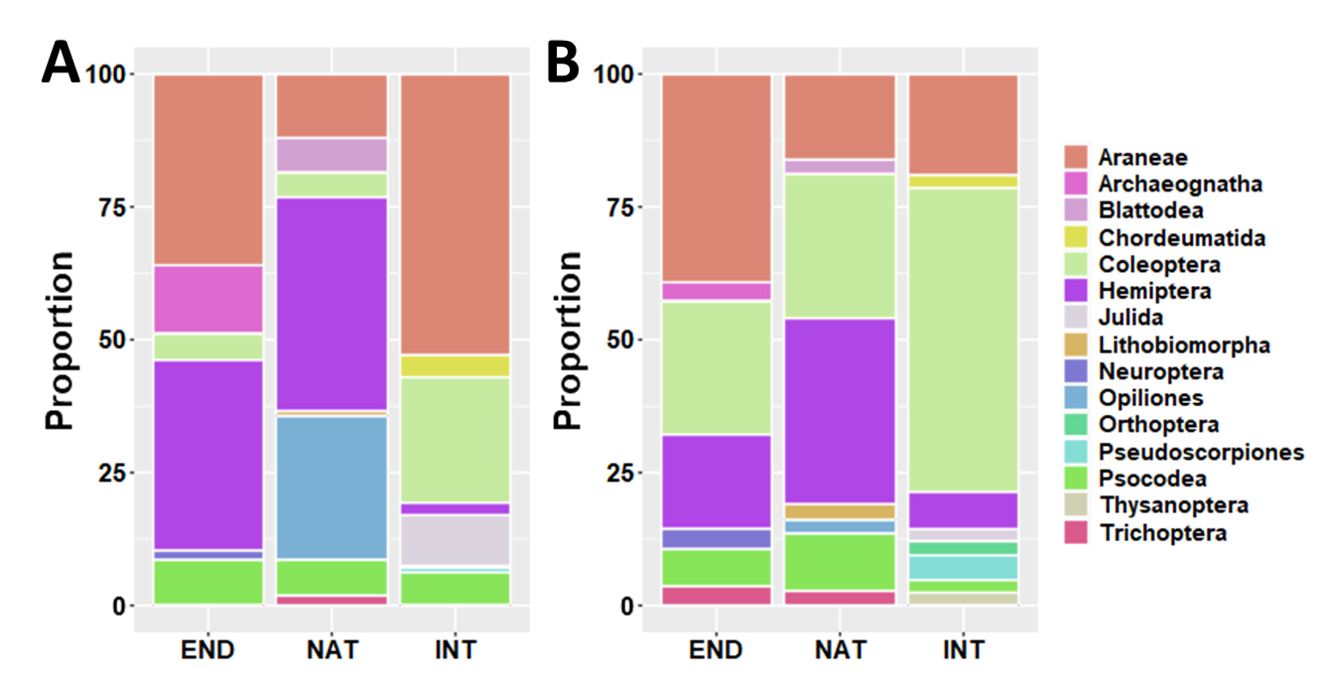

Fig. A.1. Arthropods orders sampled with SLAM in the three habitats (EDGE, CENTER, DEEP) classified within their biogeographical categories (END - endemic species, NAT - native non-endemic species and INT - introduced species). Overall proportion of individuals (A) and proportion of species (B). The graphic should be displayed in color to differentiate the fifteen orders.

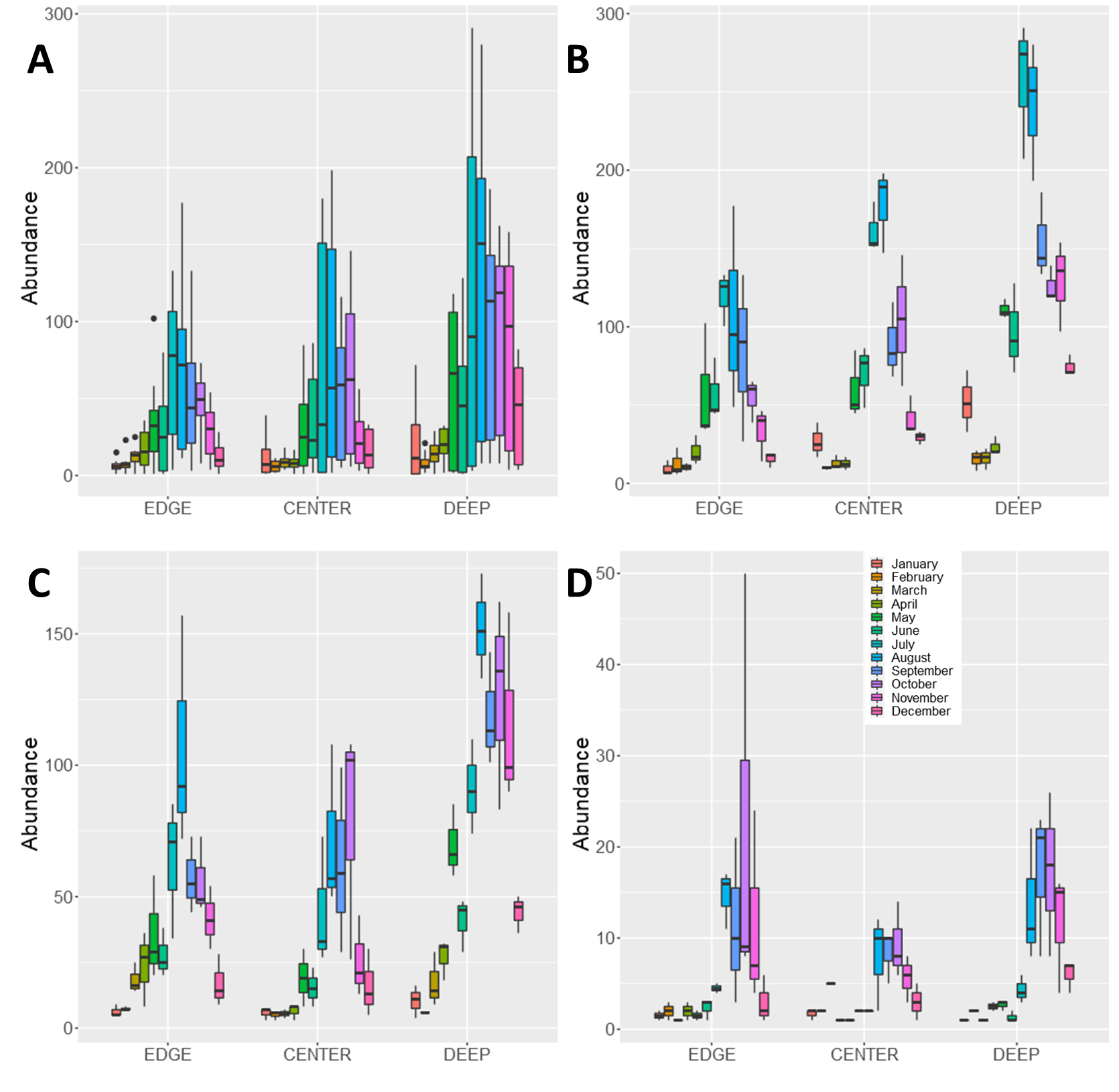

Fig. A.2. Arthropod abundance sampled per month in three habitats (forest EDGE, CENTER and DEEP) of Terra-Brava native forest fragment. Boxplots present abundance data for all species A), and for the three biogeographic groups: endemic B), native non-endemic, C) and introduced species D).

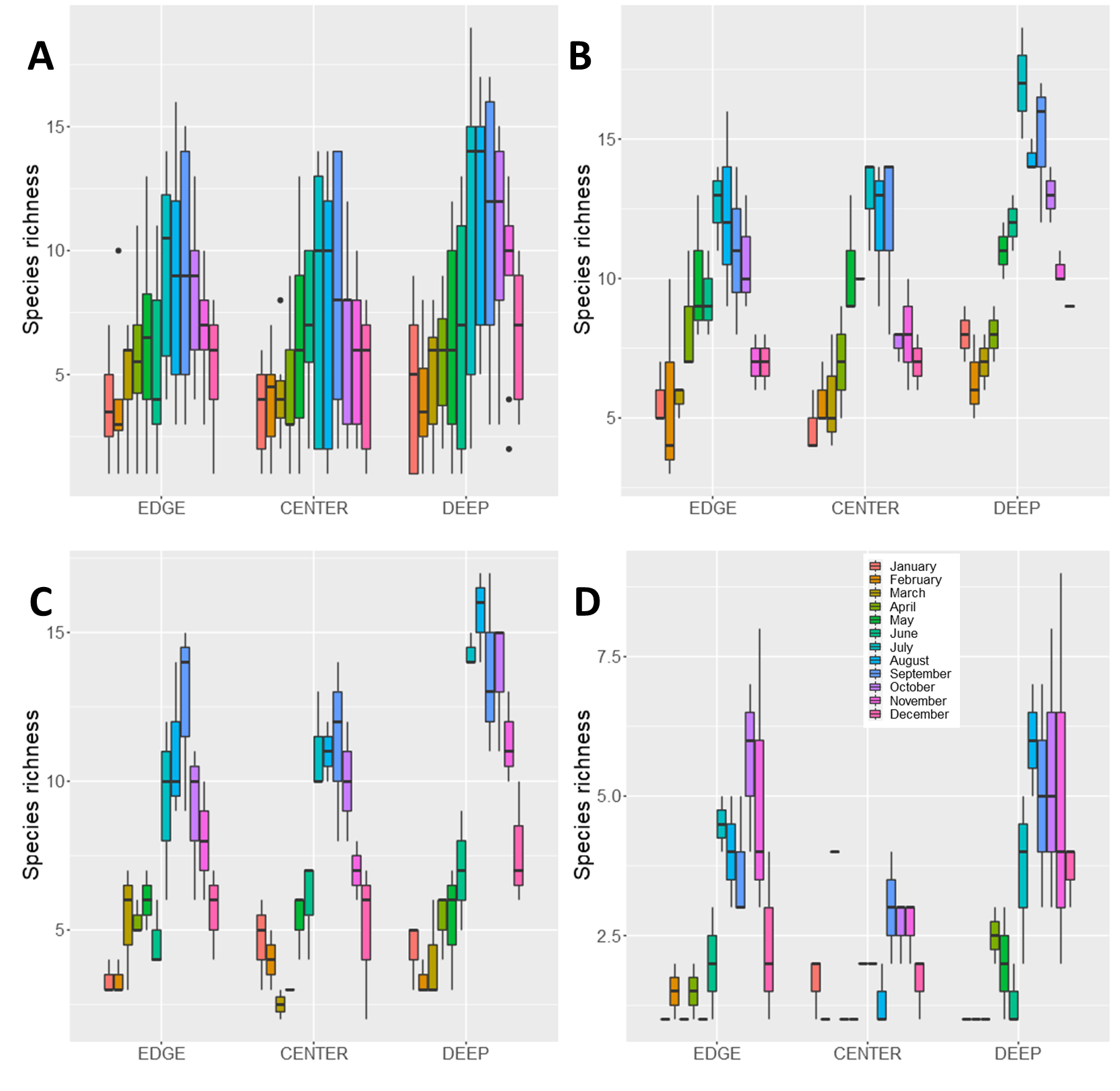

Fig. A.3. Arthropod species richness (chao1, q0) sampled per month in three habitats (forest EDGE, CENTER and DEEP) of Terra-Brava native forest fragment. Boxplots present species richness data for all species A), and for the three biogeographic groups: endemic B), native non-endemic, C) and introduced species D).

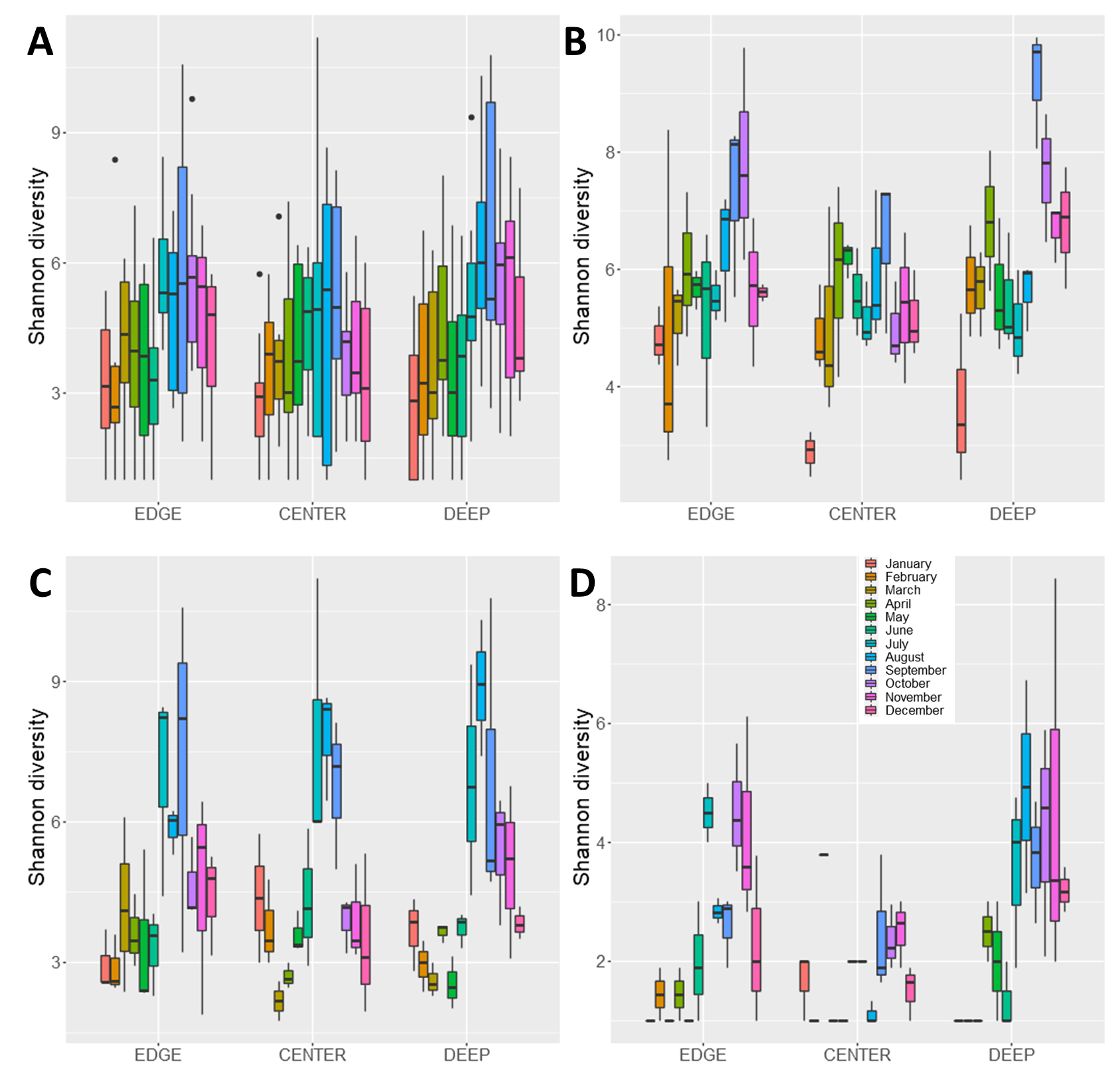

Fig. A.4. Arthropod species diversity (Shannon diversity, q1) sampled per month in three habitats (forest EDGE, CENTER and DEEP) of Terra-Brava native forest fragment. Boxplots present Shannon diversity data for all species A), and for the three biogeographic groups: endemic B), native non-endemic, C) and introduced species D).

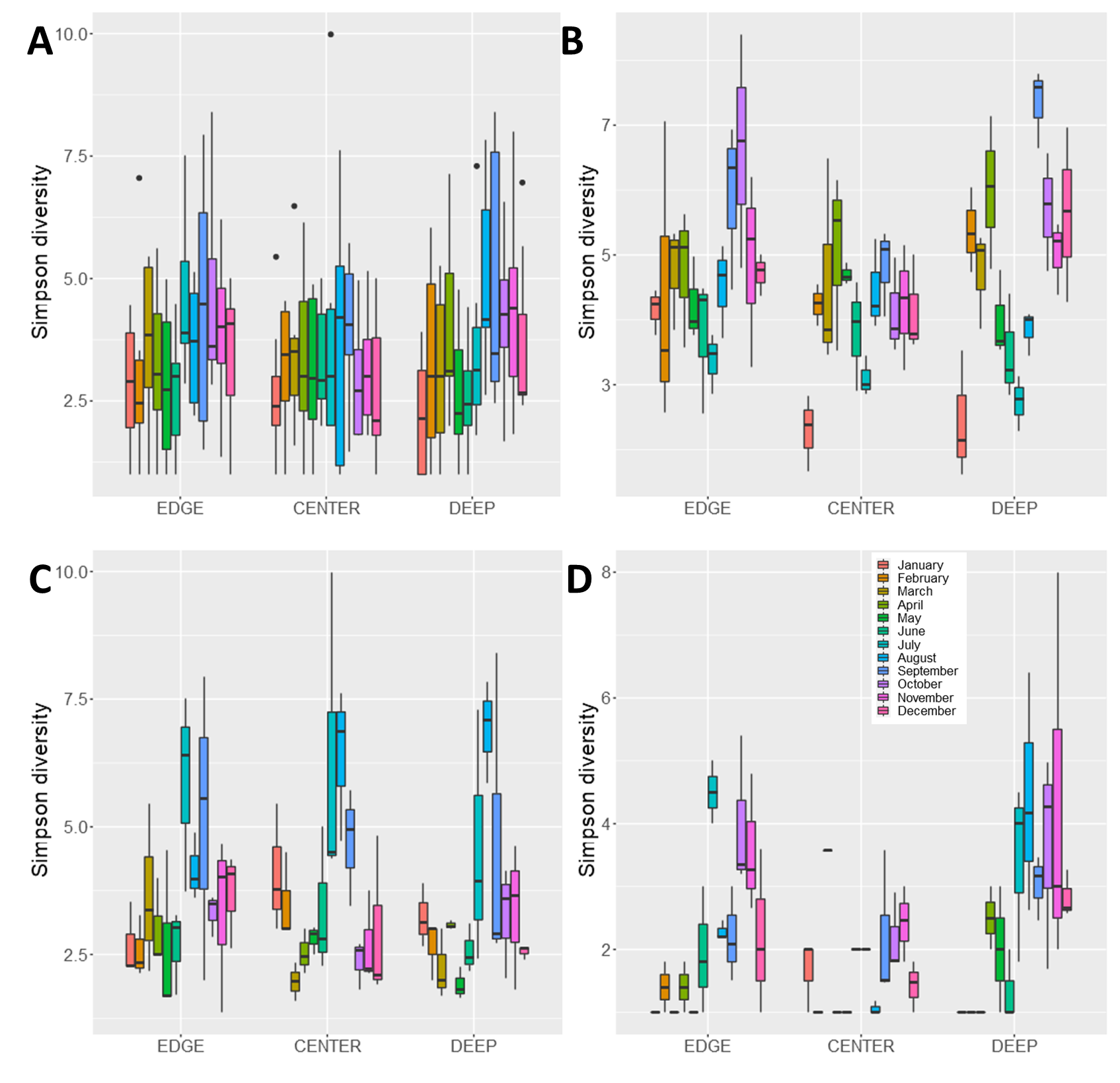

Fig. A.5. Arthropod species diversity (Simpson diversity, q2) sampled per month in three habitats (forest EDGE, CENTER and DEEP) of Terra-Brava native forest fragment. Boxplots present Simpson diversity data for all species A), and for the three biogeographic groups: endemic B), native non-endemic, C) and introduced species D).

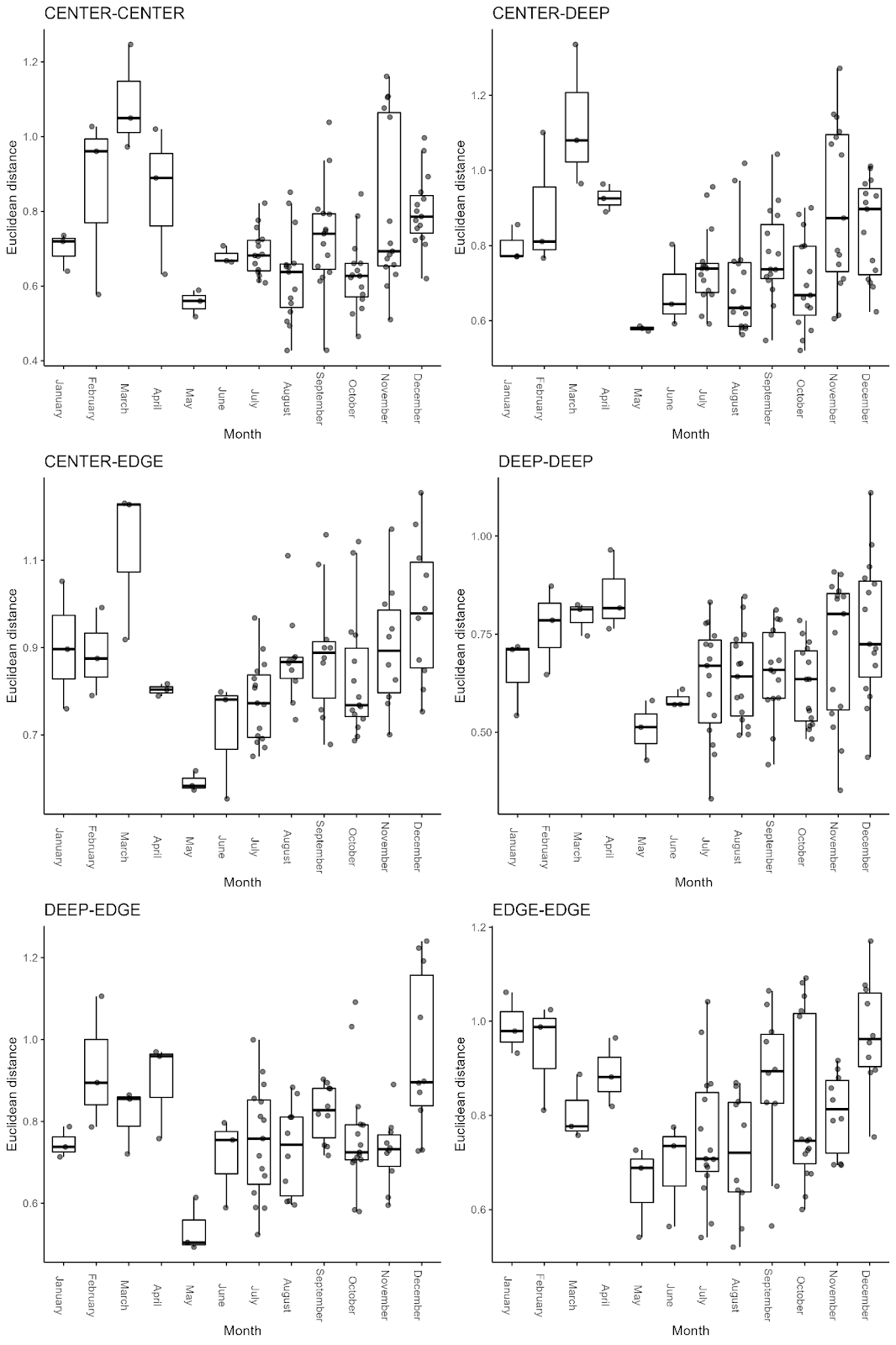

Fig. A.6. Euclidean distances comparing within-habitat and between-habitat arthropods sampled per month in Terra-Brava native forest fragment. Boxplots present first quartile, median, and third quartile, and outlier values.

#### Tables

Table. A.1. Total arthropod abundance and richness (and their proportions) sampled SLAM in the three habitats (EDGE, CENTER, DEEP) of Terra-Brava native forest fragment. Overall species (All species) and groups classified within their biogeographical categories (endemic species, native non-endemic species and introduced species).

|  | ***Number of individuals*** | ***Number of species*** | ***Proportion of number of individuals (%)*** | ***Proportion of number of species (%)*** |
| --- | --- | --- | --- | --- |
| **All species** | 13516 | 107 | 100 | 100 |
| **Endemic** | 8026 | 28 | 60 | 26 |
| **Native** | 4906 | 37 | 36 | 35 |
| **Introduced** | 584 | 42 | 4 | 39 |

Table A.2. ANOVA tables of mixed effect models of total arthropods sampled per month in three habitats (forest EDGE, CENTER and DEEP) of Terra-Brava native forest fragment. Models for all species and for species groups classified within their biogeographical categories (endemic species, native non-endemic species and Introduced species). Total abundance is the dependent variable and the three habitats, the explanatory variables. Pairwise comparisons showed significant high abundance in EDGE than in DEEP and in DEEP than in CENTER whereas not difference was found between EDGE and CENTER habitats for models with all species, endemic, native non-endemic species. Abundance of introduced species did not significantly differ between the three habitats.

|  | ***numDEF*** | ***denDF*** | ***F-value*** | ***P*** |
| --- | --- | --- | --- | --- |
| **All species** |  |  |  |  |
| **Intercept** | 1 | 6 | 362.27 | <0.001 |
| **Habitat** | 2 | 6 | 19.43 | <0.010 |
| **Endemic** |  |  |  |  |
| **Intercept** | 1 | 6 | 356.32 | <0.001 |
| **Habitat** | 2 | 6 | 20.12 | <0.010 |
| **Native** |  |  |  |  |
| **Intercept** | 1 | 6 | 143.80 | <0.001 |
| **Habitat** | 2 | 6 | 9.63 | <0.050 |
| **Introduced** |  |  |  |  |
| **Intercept** | 1 | 6 | 33.88 | <0.010 |
| **Habitat** | 2 | 6 | 1.01 | 0.418 |

Table A.3. ANOVA tables of mixed effect models of total arthropods sampled per month in three habitats (forest EDGE, CENTER and DEEP) of Terra-Brava native forest fragment. Models for all species and for species groups classified within their biogeographical categories (endemic species, native non-endemic species and Introduced species). Species richness (chao1, q0) is the dependent variable and the three habitats, the explanatory variables. Pairwise comparisons confirmed no significant difference of species richness between the three habitats for the four models.

|  | ***numDEF*** | ***denDF*** | ***F-value*** | ***P*** |
| --- | --- | --- | --- | --- |
| **All species** |  |  |  |  |
| **Intercept** | 1 | 18 | 461.351 | <0.0001 |
| **Habitat** | 2 | 6 | 1.456 | 0.305 |
| **Endemic** |  |  |  |  |
| **Intercept** | 1 | 6 | 2088.643 | <0.0001 |
| **Habitat** | 2 | 6 | 2.786 | 0.139 |
| **Native** |  |  |  |  |
| **Intercept** | 1 | 6 | 871.244 | <0.0001 |
| **Habitat** | 2 | 6 | 3.146 | 0.116 |
| **Introduced** |  |  |  |  |
| **Intercept** | 1 | 6 | 178.513 | <0.0001 |
| **Habitat** | 2 | 6 | 3.744 | 0.088 |

Table A.4. ANOVA tables of mixed effect models of total arthropods sampled per month in three habitats (forest EDGE, CENTER and DEEP) of Terra-Brava native forest fragment. Models for all species and for species groups classified within their biogeographical categories (endemic species, native non-endemic species and Introduced species). Species diversity (Shannon diversity, q1) is the dependent variable and the three habitats, the explanatory variables. Pairwise comparisons confirmed no significant difference of diversity values between the three habitats for the four models.

|  | ***numDEF*** | ***denDF*** | ***F-value*** | ***P*** |
| --- | --- | --- | --- | --- |
| **All species** |  |  |  |  |
| **Intercept** | 1 | 18 | 340.249 | <0.0001 |
| **Habitat** | 2 | 6 | 2.676 | 0.148 |
| **Endemic** |  |  |  |  |
| **Intercept** | 1 | 6 | 1193.767 | <0.0001 |
| **Habitat** | 2 | 6 | 5.131 | 0.050 |
| **Native** |  |  |  |  |
| **Intercept** | 1 | 6 | 174.040 | <0.0001 |
| **Habitat** | 2 | 6 | 0.008 | 0.992 |
| **Introduced** |  |  |  |  |
| **Intercept** | 1 | 6 | 68.095 | <0.001 |
| **Habitat** | 2 | 6 | 3.556 | 0.096 |

Table A.5. ANOVA tables of mixed effect models of total arthropods sampled per month in three habitats (forest EDGE, CENTER and DEEP) of Terra-Brava native forest fragment. Models for all species and for species groups classified within their biogeographical categories (endemic species, native non-endemic species and Introduced species). Species diversity (Simpson diversity, q2) is the dependent variable and the three habitats, the explanatory variables. Pairwise comparisons confirmed no significant difference of diversity values between the three habitats for the four models.

|  | ***numDEF*** | ***denDF*** | ***F-value*** | ***P*** |
| --- | --- | --- | --- | --- |
| **All species** |  |  |  |  |
| **Intercept** | 1 | 18 | 214.029 | <0.0001 |
| **Habitat** | 2 | 6 | 2.638 | 0.151 |
| **Endemic** |  |  |  |  |
| **Intercept** | 1 | 6 | 856.840 | <0.0001 |
| **Habitat** | 2 | 6 | 3.274 | 0.109 |
| **Native** |  |  |  |  |
| **Intercept** | 1 | 6 | 91.125 | <0.0001 |
| **Habitat** | 2 | 6 | 0.119 | 0.890 |
| **Introduced** |  |  |  |  |
| **Intercept** | 1 | 6 | 45.624 | <0.01 |
| **Habitat** | 2 | 6 | 3.398 | 0.103 |

Table A.6. Parameters of partial dbRDA models, in which sampling year and month were partialled out (partial dbRDA), run on all species, and for the three biogeographic categories (endemic species, native non-endemic species and Introduced species). Arthropods sampled per month in three habitats (forest EDGE, CENTER and DEEP) of Terra-Brava native forest fragment.

|  | ***DF*** | ***Sum of Squares*** | ***F-value*** | ***P*** |
| --- | --- | --- | --- | --- |
| **All species** |  |  |  |  |
| **Habitat** | 2 | 5.107 | 8.823 | 0.001 |
| **Residual** | 143 | 41.384 |  |  |
| **Endemic** |  |  |  |  |
| **Habitat** | 2 | 3.499 | 5.397 | 0.001 |
| **Residual** | 143 | 46.356 |  |  |
| **Native** |  |  |  |  |
| **Habitat** | 2 | 2.147 | 2.712 | 0.001 |
| **Residual** | 142 | 56.21 |  |  |
| **Introduced** |  |  |  |  |
| **Habitat** | 2 | 4.048 | 3.809 | 0.001 |
| **Residual** | 112 | 59.51 |  |  |

Table A.7. Parameters of the standard dbRDA models, in which sampling year and month were included as interacting factors, run on all species, and for the three biogeographic categories (endemic species, native non-endemic species and Introduced species). Arthropods sampled per month in three habitats (forest EDGE, CENTER and DEEP) of Terra-Brava native forest fragment.

|  | ***DF*** | ***Sum of Squares*** | ***F-value*** | ***P*** |
| --- | --- | --- | --- | --- |
| **All species** |  |  |  |  |
| **Habitat** | 2 | 5.1372 | 9.6179 | 0.001 |
| **Year** | 1 | 1.7501 | 6.5529 | 0.001 |
| **Month** | 11 | 25.9913 | 8.8474 | 0.001 |
| **Habitat × Year** | 2 | 0.9432 | 1.7658 | 0.019 |
| **Habitat × Month** | 22 | 7.9727 | 1.3569 | 0.001 |
| **Year × Month** | 5 | 2.3885 | 1.7887 | 0.002 |
| **Habitat × year × Month** | 10 | 2.3047 | 0.863 | 0.848 |
| **Residual** | 104 | 27.7749 |  |  |
| **Endemic** |  |  |  |  |
| **Habitat** | 2 | 5.031 | 12.143 | 0.001 |
| **Year** | 1 | 1.371 | 6.620 | 0.001 |
| **Month** | 11 | 23.116 | 10.145 | 0.001 |
| **Habitat × Year** | 2 | 0.899 | 2.169 | 0.008 |
| **Habitat × Month** | 22 | 6.581 | 1.444 | 0.003 |
| **Year × Month** | 5 | 1.421 | 1.372 | 0.072 |
| **Habitat × year × Month** | 10 | 1.898 | 0.916 | 0.7 |
| **Residual** | 104 | 21.543 |  |  |
| **Native** |  |  |  |  |
| **Habitat** | 2 | 3.375 | 5.374 | 0.001 |
| **Year** | 1 | 2.141 | 6.818 | 0.001 |
| **Month** | 11 | 28.222 | 8.169 | 0.001 |
| **Habitat × Year** | 2 | 1.059 | 1.685 | 0.048 |
| **Habitat × Month** | 22 | 8.854 | 1.281 | 0.015 |
| **Year x Month** | 5 | 3.448 | 2.196 | 0.001 |
| **Habitat × year × Month** | 10 | 2.627 | 0.837 | 0.842 |
| **Residual** | 103 | 32.349 |  |  |
| **Introduced** |  |  |  |  |
| **Habitat** | 2 | 7.078 | 5.990 | 0.001 |
| **Year** | 1 | 0.949 | 1.606 | 0.096 |
| **Month** | 11 | 12.795 | 1.969 | 0.001 |
| **Habitat × Year** | 2 | 1.466 | 1.241 | 0.189 |
| **Habitat × Month** | 22 | 19.031 | 1.464 | 0.001 |
| **Year × Month** | 5 | 3.187 | 1.079 | 0.332 |
| **Habitat × year × Month** | 9 | 4.089 | 0.769 | 0.951 |
| **Residual** | 74 | 43.719 |  |  |

Table A.8. Summary of the GLMM of total arthropods abundance for endemic species. Species were sampled in the native forest fragment Terra-Brava in three habitats from border (EDGE) to the core forest (DEEP) and in an intermediate habitat (CENTER). Variance explained by fixed effects: Marginal R^2^ = 0.91. Variance explained by the entire model: Conditional R^2^ = 0.93.

|  | ***Estimate*** | ***Std. Error*** | ***z value*** | ***P*** |
| --- | --- | --- | --- | --- |
| **Habitat** | |  |  |  |
| Intercept | 2.254 | 0.198 | 11.366 | <0.0001 |
| Center | 1.039 | 0.238 | 4.370 | <0.0001 |
| Deep | 1.695 | 0.225 | 7.538 | <0.0001 |
| **Month** | | |  |  |
| February | 0.270 | 0.247 | 1.096 | 0.273 |
| March | 0.067 | 0.258 | 0.258 | 0.796 |
| April | 0.744 | 0.226 | 3.297 | <0.001 |
| May | 1.792 | 0.201 | 8.933 | <0.0001 |
| June | 1.780 | 0.201 | 8.868 | <0.0001 |
| July | 1.823 | 0.193 | 9.443 | <0.0001 |
| August | 1.851 | 0.194 | 9.541 | <0.0001 |
| September | 1.601 | 0.196 | 8.158 | <0.0001 |
| October | 1.039 | 0.201 | 5.160 | <0.0001 |
| November | 0.685 | 0.211 | 3.246 | 0.001 |
| December | -0.092 | 0.237 | -0.386 | 0.699 |
| **Habitat × Month** | | | |  |
| Center*February | -1.264 | 0.326 | -3.872 | <0.001 |
| Center*March | -0.798 | 0.324 | -2.465 | 0.014 |
| Center*April | -1.500 | 0.299 | -5.014 | <0.0001 |
| Center*May | -0.993 | 0.241 | -4.120 | <0.0001 |
| Center*June | -0.823 | 0.240 | -3.435 | <0.001 |
| Center*July | -0.728 | 0.227 | -3.204 | 0.001 |
| Center*August | -0.658 | 0.228 | -2.891 | 0.004 |
| Center*September | -1.102 | 0.234 | -4.713 | <0.0001 |
| Center*October | -0.381 | 0.237 | -1.608 | 0.108 |
| Center*November | -0.944 | 0.255 | -3.707 | <0.001 |
| Center*December | -0.519 | 0.283 | -1.834 | 0.067 |
| Deep*February | -1.492 | 0.298 | -5.001 | <0.0001 |
| Deep*March | -1.245 | 0.307 | -4.062 | <0.0001 |
| Deep*April | -1.545 | 0.268 | -5.775 | <0.0001 |
| Deep*May | -1.033 | 0.223 | -4.638 | <0.0001 |
| Deep*June | -1.160 | 0.224 | -5.180 | <0.0001 |
| Deep*July | -0.917 | 0.212 | -4.324 | <0.0001 |
| Deep*August | -1.009 | 0.213 | -4.736 | <0.0001 |
| Deep*September | -1.204 | 0.217 | -5.550 | <0.0001 |
| Deep*October | -0.848 | 0.223 | -3.804 | <0.001 |
| Deep*November | -0.469 | 0.231 | -2.029 | 0.042 |
| Deep*December | -0.244 | 0.259 | -0.942 | 0.346 |

Table A.9. Summary of the GLMM of total arthropods abundance for native species. Species were sampled in the native forest fragment Terra-Brava in three habitats from border (EDGE) to the core forest (DEEP) and in an intermediate habitat (CENTER). Variance explained by fixed effects: Marginal R^2^ = 0.95. Variance explained by the entire model: Conditional R^2^ = 0.97.

|  | ***Estimate*** | ***Std. Error*** | ***z value*** | ***P*** |
| --- | --- | --- | --- | --- |
| **Habitat** | |  |  |  |
| Intercept | 1.817 | 0.284 | 6.404 | <0.0001 |
| Center | -0.165 | 0.409 | -0.404 | 0.686 |
| Deep | 0.511 | 0.375 | 1.362 | 0.173 |
| **Month** | | |  |  |
| February | 0.147 | 0.313 | 0.468 | 0.63972 |
| March | 1.063 | 0.266 | 3.994 | <0.0001 |
| April | 1.318 | 0.258 | 5.104 | <0.0001 |
| May | 1.728 | 0.249 | 6.943 | <0.0001 |
| June | 1.474 | 0.254 | 5.797 | <0.0001 |
| July | 1.609 | 0.241 | 6.689 | <0.0001 |
| August | 2.355 | 0.236 | 9.968 | <0.0001 |
| September | 1.731 | 0.242 | 7.156 | <0.0001 |
| October | 1.486 | 0.242 | 6.141 | <0.0001 |
| November | 1.412 | 0.246 | 5.731 | <0.0001 |
| December | 0.516 | 0.269 | 1.917 | 0.055 |
| **Habitat × Month** |  |  |  |  |
| Center*February | -0.272 | 0.473 | -0.575 | 0.565 |
| Center*March | -1.301 | 0.470 | -2.770 | 0.006 |
| Center*April | -1.207 | 0.422 | -2.860 | 0.004 |
| Center*May | -0.519 | 0.372 | -1.394 | 0.163 |
| Center*June | -0.479 | 0.381 | -1.257 | 0.209 |
| Center*July | -0.245 | 0.352 | -0.696 | 0.486 |
| Center*August | -0.511 | 0.345 | -1.479 | 0.139 |
| Center*September | -0.026 | 0.350 | -0.076 | 0.940 |
| Center*October | 0.451 | 0.349 | 1.293 | 0.196 |
| Center*November | -0.595 | 0.364 | -1.633 | 0.102 |
| Center*December | -0.171 | 0.390 | -0.438 | 0.662 |
| Deep*February | -0.690 | 0.431 | -1.601 | 0.109 |
| Deep*March | -0.546 | 0.350 | -1.560 | 0.119 |
| Deep*April | -0.358 | 0.334 | -1.072 | 0.284 |
| Deep*May | 0.180 | 0.315 | 0.572 | 0.567 |
| Deep*June | -0.104 | 0.324 | -0.322 | 0.748 |
| Deep*July | -0.123 | 0.306 | -0.403 | 0.687 |
| Deep*August | -0.358 | 0.300 | -1.190 | 0.234 |
| Deep*September | 0.019 | 0.306 | 0.063 | 0.949 |
| Deep*October | 0.329 | 0.306 | 1.077 | 0.281 |
| Deep*November | 0.310 | 0.310 | 1.002 | 0.316 |
| Deep*December | 0.240 | 0.335 | 0.717 | 0.473 |

Table A.10. Summary of the GLMM of total arthropods abundance for introduced species. Species were sampled in the native forest fragment Terra-Brava in three habitats from border (EDGE) to the core forest (DEEP) and in an intermediate habitat (CENTER). Variance explained by fixed effects: Marginal R^2^ = 0.57. Variance explained by the entire model: Conditional R^2^ = 0.70.

|  | ***Estimate*** | ***Std. Error*** | ***z value*** | ***P*** |
| --- | --- | --- | --- | --- |
| **Habitat** |  |  |  |  |
| Intercept | 0.633 | 0.611 | 1.036 | 0.300 |
| Center | -0.144 | 0.782 | -0.184 | 0.854 |
| Deep | -0.677 | 0.863 | -0.784 | 0.433 |
| **Month** |  |  |  |  |
| February | -0.244 | 0.770 | -0.317 | 0.751 |
| March | -1.173 | 1.164 | -1.008 | 0.313 |
| April | 0.288 | 0.764 | 0.377 | 0.706 |
| May | -0.374 | 0.824 | -0.453 | 0.650 |
| June | 0.116 | 0.694 | 0.167 | 0.867 |
| July | 0.405 | 0.667 | 0.608 | 0.543 |
| August | 1.424 | 0.603 | 2.361 | 0.018 |
| September | 1.279 | 0.610 | 2.096 | 0.036 |
| October | 1.682 | 0.595 | 2.827 | 0.005 |
| November | 1.195 | 0.608 | 1.966 | 0.049 |
| December | 0.367 | 0.671 | 0.548 | 0.584 |
| **Habitat × Month** |  |  |  |  |
| Center*February | 0.398 | 1.022 | 0.389 | 0.697 |
| Center*March | 2.080 | 1.328 | 1.566 | 0.117 |
| Center*April | -0.633 | 1.342 | -0.472 | 0.637 |
| Center*May | -0.025 | 1.177 | -0.022 | 0.983 |
| Center*June | 0.127 | 1.094 | 0.116 | 0.908 |
| Center*July | -0.223 | 0.901 | -0.248 | 0.804 |
| Center*August | -0.396 | 0.778 | -0.509 | 0.611 |
| Center*September | -0.362 | 0.783 | -0.463 | 0.643 |
| Center*October | -0.652 | 0.768 | -0.849 | 0.396 |
| Center*November | -0.665 | 0.793 | -0.838 | 0.402 |
| Center*December | -0.028 | 0.873 | -0.033 | 0.974 |
| Deep*February | 0.774 | 1.084 | 0.714 | 0.475 |
| Deep*March | 1.015 | 1.641 | 0.618 | 0.536 |
| Deep*April | 0.465 | 1.057 | 0.440 | 0.660 |
| Deep*May | 1.354 | 1.066 | 1.270 | 0.205 |
| Deep*June | 0.172 | 1.032 | 0.166 | 0.868 |
| Deep*July | 0.368 | 0.925 | 0.398 | 0.691 |
| Deep*August | 0.498 | 0.850 | 0.586 | 0.558 |
| Deep*September | 0.881 | 0.851 | 1.034 | 0.301 |
| Deep*October | 0.478 | 0.840 | 0.568 | 0.570 |
| Deep*November | 0.682 | 0.855 | 0.797 | 0.425 |
| Deep*December | 0.845 | 0.916 | 0.922 | 0.356 |

Table A.11. Summary of the GLMM of arthropods species richness (Chao1, q0) for endemic species. Species were sampled in the native forest fragment Terra-Brava in three habitats from border (EDGE) to the core forest (DEEP) and in an intermediate habitat (CENTER). Variance explained by fixed effects: Marginal R^2^ = 0.49. Variance explained by the entire model: Conditional R^2^ = 0.49.

|  | ***Estimate*** | ***Std. Error*** | ***z value*** | ***P*** |
| --- | --- | --- | --- | --- |
| **Habitat** |  |  |  |  |
| Intercept | 1.74E+00 | 2.43E-01 | 7.152 | <0.0001 |
| Center | -1.94E-01 | 3.61E-01 | -0.538 | 0.591 |
| Deep | 3.45E-01 | 3.17E-01 | 1.088 | 0.277 |
| **Month** |  |  |  |  |
| February | 1.17E-05 | 3.43E-01 | 0 | 0.999 |
| March | 1.10E-05 | 3.43E-01 | 0 | 0.999 |
| April | 3.86E-01 | 3.14E-01 | 1.227 | 0.220 |
| May | 5.68E-01 | 3.04E-01 | 1.871 | 0.061 |
| June | 4.99E-01 | 3.08E-01 | 1.623 | 0.105 |
| July | 8.04E-01 | 2.92E-01 | 2.757 | 0.006 |
| August | 7.78E-01 | 2.93E-01 | 2.654 | 0.008 |
| September | 6.63E-01 | 2.99E-01 | 2.222 | 0.026 |
| October | 6.33E-01 | 3.00E-01 | 2.108 | 0.035 |
| November | 2.11E-01 | 3.26E-01 | 0.648 | 0.517 |
| December | 2.11E-01 | 3.26E-01 | 0.648 | 0.517 |
| **Habitat × Month** | |  |  |  |
| Center*February | 1.94E-01 | 4.98E-01 | 0.39 | 0.697 |
| Center*March | 1.94E-01 | 4.98E-01 | 0.39 | 0.697 |
| Center*April | 1.98E-02 | 4.67E-01 | 0.042 | 0.966 |
| Center*May | 2.27E-01 | 4.43E-01 | 0.513 | 0.608 |
| Center*June | 2.63E-01 | 4.46E-01 | 0.589 | 0.556 |
| Center*July | 2.20E-01 | 4.27E-01 | 0.516 | 0.606 |
| Center*August | 1.67E-01 | 4.30E-01 | 0.388 | 0.698 |
| Center*September | 2.81E-01 | 4.34E-01 | 0.648 | 0.517 |
| Center*October | -1.36E-01 | 4.53E-01 | -0.301 | 0.764 |
| Center*November | 3.28E-01 | 4.69E-01 | 0.699 | 0.484 |
| Center*December | 1.94E-01 | 4.75E-01 | 0.409 | 0.683 |
| Deep*February | -2.34E-01 | 4.60E-01 | -0.507 | 0.612 |
| Deep*March | -1.34E-01 | 4.55E-01 | -0.294 | 0.769 |
| Deep*April | -3.86E-01 | 4.27E-01 | -0.904 | 0.366 |
| Deep*May | -2.50E-01 | 4.05E-01 | -0.616 | 0.538 |
| Deep*June | -9.35E-02 | 4.05E-01 | -0.231 | 0.817 |
| Deep*July | -5.06E-02 | 3.83E-01 | -0.132 | 0.895 |
| Deep*August | -1.95E-01 | 3.88E-01 | -0.501 | 0.616 |
| Deep*September | -3.47E-02 | 3.91E-01 | -0.089 | 0.929 |
| Deep*October | -1.47E-01 | 3.97E-01 | -0.371 | 0.711 |
| Deep*November | 4.46E-02 | 4.25E-01 | 0.105 | 0.916 |
| Deep*December | -9.35E-02 | 4.30E-01 | -0.217 | 0.828 |

Table A.12. Summary of the GLMM of arthropods species richness (Chao1, q0) for native species. Species were sampled in the native forest fragment Terra-Brava in three habitats from border (EDGE) to the core forest (DEEP) and in an intermediate habitat (CENTER). Variance explained by fixed effects: Marginal R^2^ = 0.63. Variance explained by the entire model: Conditional R^2^ = 0.63.

|  | ***Estimate*** | ***Std. Error*** | ***z value*** | ***P*** |
| --- | --- | --- | --- | --- |
| **Habitat** |  |  |  |  |
| Intercept | 1.20E+00 | 3.16E-01 | 3.807 | <0.001 |
| Center | 3.37E-01 | 4.14E-01 | 0.813 | 0.416 |
| Deep | 2.62E-01 | 4.21E-01 | 0.624 | 0.533 |
| **Month** |  |  |  |  |
| February | 5.05E-07 | 4.47E-01 | <0.0001 | 0.999 |
| March | 4.70E-01 | 4.03E-01 | 1.166 | 0.244 |
| April | 4.70E-01 | 4.03E-01 | 1.166 | 0.244 |
| May | 5.88E-01 | 3.94E-01 | 1.49 | 0.136 |
| June | 3.37E-01 | 4.14E-01 | 0.813 | 0.416 |
| July | 1.03E+00 | 3.68E-01 | 2.795 | 0.005 |
| August | 1.19E+00 | 3.61E-01 | 3.307 | <0.001 |
| September | 1.34E+00 | 3.55E-01 | 3.756 | <0.001 |
| October | 9.93E-01 | 3.70E-01 | 2.683 | 0.007 |
| November | 8.76E-01 | 3.76E-01 | 2.326 | 0.020 |
| December | 5.31E-01 | 3.99E-01 | 1.331 | 0.183 |
| **Habitat × Month** |  |  |  |  |
| Center*February | -1.54E-01 | 5.96E-01 | -0.259 | 0.796 |
| Center*March | -1.09E+00 | 6.59E-01 | -1.661 | 0.097 |
| Center*April | -9.12E-01 | 5.87E-01 | -1.552 | 0.121 |
| Center*May | -4.54E-01 | 5.38E-01 | -0.844 | 0.398 |
| Center*June | -8.52E-02 | 5.46E-01 | -0.156 | 0.876 |
| Center*July | -1.72E-01 | 4.87E-01 | -0.353 | 0.724 |
| Center*August | -3.37E-01 | 4.82E-01 | -0.699 | 0.485 |
| Center*September | -4.48E-01 | 4.77E-01 | -0.939 | 0.348 |
| Center*October | -2.31E-01 | 4.92E-01 | -0.470 | 0.638 |
| Center*November | -4.70E-01 | 5.11E-01 | -0.920 | 0.357 |
| Center*December | -4.62E-01 | 5.45E-01 | -0.847 | 0.397 |
| Deep*February | -2.62E-01 | 6.14E-01 | -0.427 | 0.669 |
| Deep*March | -5.50E-01 | 5.68E-01 | -0.968 | 0.333 |
| Deep*April | -2.62E-01 | 5.50E-01 | -0.477 | 0.633 |
| Deep*May | -3.80E-01 | 5.43E-01 | -0.700 | 0.484 |
| Deep*June | 1.43E-01 | 5.44E-01 | 0.263 | 0.793 |
| Deep*July | 1.67E-01 | 4.86E-01 | 0.343 | 0.732 |
| Deep*August | 9.13E-02 | 4.78E-01 | 0.191 | 0.849 |
| Deep*September | -1.86E-01 | 4.77E-01 | -0.391 | 0.696 |
| Deep*October | 1.55E-01 | 4.88E-01 | 0.318 | 0.750 |
| Deep*November | 8.59E-02 | 4.98E-01 | 0.173 | 0.863 |
| Deep*December | 3.99E-02 | 5.28E-01 | 0.076 | 0.940 |

Table A.13. Summary of the GLMM arthropods species richness (Chao1, q0) for introduced species. Species were sampled in the native forest fragment Terra-Brava in three habitats from border (EDGE) to the core forest (DEEP) and in an intermediate habitat (CENTER). Variance explained by fixed effects: Marginal R^2^ = 0.46. Variance explained by the entire model: Conditional R^2^ = 0.46.

|  | ***Estimate*** | ***Std. Error*** | ***z value*** | ***P*** |
| --- | --- | --- | --- | --- |
| **Habitat** |  |  |  |  |
| Intercept | 0.006 | 0.710 | 0.008 | 0.992 |
| Center | 0.504 | 0.840 | 0.600 | 0.548 |
| Deep | -0.007 | 0.916 | -0.007 | 0.994 |
| **Month** |  |  |  |  |
| February | 0.395 | 0.920 | 0.430 | 0.667 |
| March | -0.020 | 1.243 | -0.016 | 0.987 |
| April | 0.405 | 0.913 | 0.444 | 0.657 |
| May | -0.010 | 1.006 | -0.010 | 0.992 |
| June | 0.687 | 0.820 | 0.838 | 0.402 |
| July | 1.504 | 0.782 | 1.924 | 0.054 |
| August | 1.380 | 0.767 | 1.798 | 0.072 |
| September | 1.293 | 0.772 | 1.674 | 0.094 |
| October | 1.728 | 0.751 | 2.301 | 0.021 |
| November | 1.603 | 0.756 | 2.119 | 0.034 |
| December | 0.841 | 0.805 | 1.044 | 0.296 |
| **Habitat × Month** | |  |  |  |
| Center*February | -0.906 | 1.243 | -0.729 | 0.466 |
| Center*March | 0.895 | 1.413 | 0.633 | 0.526 |
| Center*April | -0.916 | 1.426 | -0.642 | 0.521 |
| Center*May | -0.501 | 1.308 | -0.383 | 0.702 |
| Center*June | -0.504 | 1.172 | -0.430 | 0.667 |
| Center*July | -1.322 | 0.989 | -1.337 | 0.181 |
| Center*August | -1.603 | 1.019 | -1.573 | 0.116 |
| Center*September | -0.705 | 0.952 | -0.740 | 0.459 |
| Center*October | -1.258 | 0.943 | -1.334 | 0.182 |
| Center*November | -1.133 | 0.947 | -1.196 | 0.232 |
| Center*December | -0.841 | 1.024 | -0.821 | 0.412 |
| Deep*February | -0.401 | 1.292 | -0.310 | 0.757 |
| Deep*March | 0.006 | 1.685 | 0.004 | 0.997 |
| Deep*April | 0.506 | 1.170 | 0.432 | 0.666 |
| Deep*May | 0.703 | 1.229 | 0.572 | 0.568 |
| Deep*June | -0.399 | 1.120 | -0.356 | 0.722 |
| Deep*July | -0.205 | 1.018 | -0.201 | 0.841 |
| Deep*August | 0.412 | 0.989 | 0.417 | 0.677 |
| Deep*September | 0.317 | 0.998 | 0.317 | 0.751 |
| Deep*October | -0.054 | 0.980 | -0.055 | 0.956 |
| Deep*November | 0.007 | 0.986 | 0.007 | 0.995 |
| Deep*December | 0.459 | 1.036 | 0.443 | 0.658 |

Table A.14. Summary of the GLMM of arthropods species diversity (Shannon diversity, q1) for endemic species. Species were sampled in the native forest fragment Terra-Brava in three habitats from border (EDGE) to the core forest (DEEP) and in an intermediate habitat (CENTER). Variance explained by fixed effects: Marginal R^2^ = 0.58. Variance explained by the entire model: Conditional R^2^ = 0.59.

|  | ***Estimate*** | ***Std. Error*** | ***z value*** | ***P*** |
| --- | --- | --- | --- | --- |
| **Habitat** |  |  |  |  |
| Intercept | 4.814 | 0.550 | 8.751 | <0.0001 |
| Center | -1.940 | 0.778 | -2.494 | 0.012 |
| Deep | -1.150 | 0.778 | -1.478 | 0.139 |
| **Month** |  |  |  |  |
| February | 0.131 | 0.774 | 0.169 | 0.866 |
| March | 0.343 | 0.774 | 0.443 | 0.658 |
| April | 1.216 | 0.774 | 1.571 | 0.116 |
| May | 0.865 | 0.774 | 1.117 | 0.264 |
| June | 0.371 | 0.774 | 0.480 | 0.632 |
| July | 0.714 | 0.774 | 0.923 | 0.3562 |
| August | 1.567 | 0.774 | 2.024 | 0.043 |
| September | 2.499 | 0.774 | 3.227 | 0.001 |
| October | 3.032 | 0.774 | 3.916 | <0.0001 |
| November | 0.829 | 0.774 | 1.070 | 0.284 |
| December | 0.789 | 0.774 | 1.019 | 0.308 |
| **Habitat × Month** | |  |  |  |
| Center*February | 1.885 | 1.095 | 1.721 | 0.085 |
| Center*March | 1.807 | 1.095 | 1.650 | 0.099 |
| Center*April | 1.826 | 1.095 | 1.667 | 0.095 |
| Center*May | 2.459 | 1.095 | 2.246 | 0.025 |
| Center*June | 2.322 | 1.095 | 2.121 | 0.034 |
| Center*July | 1.549 | 1.095 | 1.415 | 0.157 |
| Center*August | 1.440 | 1.095 | 1.315 | 0.188 |
| Center*September | 1.128 | 1.095 | 1.031 | 0.303 |
| Center*October | -0.934 | 1.095 | -0.853 | 0.393 |
| Center*November | 1.673 | 1.095 | 1.528 | 0.127 |
| Center*December | 1.509 | 1.095 | 1.378 | 0.168 |
| Deep*February | 1.959 | 1.095 | 1.789 | 0.074 |
| Deep*March | 1.641 | 1.095 | 1.498 | 0.134 |
| Deep*April | 1.938 | 1.095 | 1.770 | 0.077 |
| Deep*May | 1.075 | 1.095 | 0.982 | 0.326 |
| Deep*June | 1.443 | 1.095 | 1.318 | 0.188 |
| Deep*July | 0.631 | 1.095 | 0.577 | 0.564 |
| Deep*August | 0.393 | 1.095 | 0.359 | 0.719 |
| Deep*September | 3.080 | 1.095 | 2.813 | 0.005 |
| Deep*October | 0.942 | 1.095 | 0.861 | 0.389 |
| Deep*November | 2.192 | 1.095 | 2.002 | 0.045 |
| Deep*December | 2.316 | 1.095 | 2.116 | 0.034 |

Table A.15. Summary of the GLMM of arthropods species diversity (Shannon diversity, q1) for native species. Species were sampled in the native forest fragment Terra-Brava in three habitats from border (EDGE) to the core forest (DEEP) and in an intermediate habitat (CENTER). Variance explained by fixed effects: Marginal R^2^ = 0.65. Variance explained by the entire model: Conditional R^2^ = 0.66.

|  | ***Estimate*** | ***Std. Error*** | ***z value*** | ***P*** |
| --- | --- | --- | --- | --- |
| **Habitat** |  |  |  |  |
| Intercept | 2.960 | 0.729 | 4.059 | <0.0001 |
| Center | 1.411 | 1.031 | 1.368 | 0.171 |
| Deep | 0.720 | 1.031 | 0.698 | 0.485 |
| **Month** |  |  |  |  |
| February | -0.078 | 1.018 | -0.077 | 0.939 |
| March | 1.235 | 1.018 | 1.213 | 0.225 |
| April | 0.660 | 1.018 | 0.648 | 0.517 |
| May | 0.435 | 1.018 | 0.427 | 0.669 |
| June | 0.337 | 1.018 | 0.331 | 0.740 |
| July | 4.070 | 1.018 | 3.998 | <0.0001 |
| August | 2.898 | 1.018 | 2.846 | 0.004 |
| September | 4.376 | 1.018 | 4.298 | <0.0001 |
| October | 1.703 | 1.018 | 1.673 | 0.094 |
| November | 1.634 | 1.018 | 1.605 | 0.108 |
| December | 1.442 | 1.018 | 1.416 | 0.157 |
| **Habitat × Month** | |  |  |  |
| Center*February | -0.551 | 1.440 | -0.382 | 0.702 |
| Center*March | -3.459 | 1.530 | -2.261 | 0.023 |
| Center*April | -2.328 | 1.440 | -1.617 | 0.106 |
| Center*May | -1.214 | 1.440 | -0.843 | 0.399 |
| Center*June | -0.400 | 1.440 | -0.278 | 0.781 |
| Center*July | -0.702 | 1.440 | -0.487 | 0.626 |
| Center*August | 0.564 | 1.440 | 0.392 | 0.695 |
| Center*September | -1.981 | 1.440 | -1.376 | 0.169 |
| Center*October | -2.191 | 1.440 | -1.522 | 0.128 |
| Center*November | -2.088 | 1.440 | -1.450 | 0.147 |
| Center*December | -2.348 | 1.440 | -1.631 | 0.102 |
| Deep*February | -0.653 | 1.440 | -0.454 | 0.650 |
| Deep*March | -2.308 | 1.440 | -1.603 | 0.109 |
| Deep*April | -0.692 | 1.440 | -0.481 | 0.631 |
| Deep*May | -1.577 | 1.440 | -1.096 | 0.273 |
| Deep*June | -0.289 | 1.440 | -0.201 | 0.841 |
| Deep*July | -0.908 | 1.440 | -0.630 | 0.528 |
| Deep*August | 2.307 | 1.440 | 1.603 | 0.109 |
| Deep*September | -1.165 | 1.440 | -0.809 | 0.418 |
| Deep*October | 0.013 | 1.440 | 0.009 | 0.993 |
| Deep*November | -0.291 | 1.440 | -0.202 | 0.840 |
| Deep*December | -1.289 | 1.440 | -0.896 | 0.370 |

Table A.16. Summary of the GLMM arthropods species diversity (Shannon diversity, q1) for introduced species. Species were sampled in the native forest fragment Terra-Brava in three habitats from border (EDGE) to the core forest (DEEP) and in an intermediate habitat (CENTER). Variance explained by fixed effects: Marginal R^2^ = 0.67. Variance explained by the entire model: Conditional R^2^ = 0.70.

|  | ***Estimate*** | ***Std. Error*** | ***z value*** | ***P*** |
| --- | --- | --- | --- | --- |
| **Habitat** |  |  |  |  |
| Intercept | 1.118 | 0.623 | 1.795 | 0.073 |
| Center | 0.549 | 0.803 | 0.684 | 0.494 |
| Deep | -0.118 | 0.803 | -0.146 | 0.884 |
| **Month** |  |  |  |  |
| February | 0.271 | 0.861 | 0.315 | 0.753 |
| March | -0.353 | 1.084 | -0.325 | 0.745 |
| April | 0.445 | 0.842 | 0.529 | 0.597 |
| May | -0.179 | 0.861 | -0.208 | 0.835 |
| June | 0.846 | 0.777 | 1.089 | 0.276 |
| July | 3.500 | 0.842 | 4.157 | <0.0001 |
| August | 1.722 | 0.777 | 2.218 | 0.027 |
| September | 1.478 | 0.777 | 1.903 | 0.057 |
| October | 3.400 | 0.777 | 4.378 | <0.0001 |
| November | 3.062 | 0.777 | 3.942 | <0.0001 |
| December | 1.142 | 0.777 | 1.471 | 0.141 |
| **Habitat × Month** | |  |  |  |
| Center*February | -0.931 | 1.157 | -0.805 | 0.421 |
| Center*March | 2.507 | 1.468 | 1.708 | 0.088 |
| Center*April | -1.131 | 1.295 | -0.873 | 0.383 |
| Center*May | -0.504 | 1.156 | -0.436 | 0.663 |
| Center*June | -0.526 | 1.254 | -0.420 | 0.675 |
| Center*July | -3.167 | 1.087 | -2.913 | 0.004 |
| Center*August | -2.278 | 1.037 | -2.197 | 0.028 |
| Center*September | -0.701 | 1.037 | -0.676 | 0.499 |
| Center*October | -2.714 | 1.037 | -2.617 | 0.009 |
| Center*November | -2.216 | 1.037 | -2.136 | 0.033 |
| Center*December | -1.296 | 1.037 | -1.250 | 0.211 |
| Deep*February | -0.375 | 1.152 | -0.326 | 0.745 |
| Deep*March | 0.012 | 1.445 | 0.008 | 0.993 |
| Deep*April | 0.951 | 1.144 | 0.831 | 0.406 |
| Deep*May | 1.179 | 1.102 | 1.070 | 0.285 |
| Deep*June | -0.512 | 1.037 | -0.494 | 0.621 |
| Deep*July | -0.949 | 1.087 | -0.873 | 0.382 |
| Deep*August | 2.209 | 1.037 | 2.130 | 0.033 |
| Deep*September | 1.241 | 1.037 | 1.196 | 0.232 |
| Deep*October | -0.211 | 1.037 | -0.203 | 0.839 |
| Deep*November | 0.537 | 1.037 | 0.518 | 0.605 |
| Deep*December | 1.052 | 1.037 | 1.014 | 0.310 |

Table A.17. Summary of the GLMM of arthropods species diversity (Simpson diversity, q2) for endemic species. Species were sampled in the native forest fragment Terra-Brava in three habitats from border (EDGE) to the core forest (DEEP) and in an intermediate habitat (CENTER). Variance explained by fixed effects: Marginal R^2^ = 0.20. Variance explained by the entire model: Conditional R^2^ = 0.20.

|  | ***Estimate*** | ***Std. Error*** | ***z value*** | ***P*** |
| --- | --- | --- | --- | --- |
| **Habitat** |  |  |  |  |
| Intercept | 1.425 | 0.283 | 5.031 | <0.0001 |
| Center | -0.595 | 0.475 | -1.253 | 0.210 |
| Deep | -0.537 | 0.466 | -1.151 | 0.250 |
| **Month** |  |  |  |  |
| February | 0.053 | 0.395 | 0.134 | 0.894 |
| March | 0.137 | 0.387 | 0.354 | 0.724 |
| April | 0.138 | 0.387 | 0.355 | 0.722 |
| May | 0.019 | 0.399 | 0.048 | 0.961 |
| June | -0.095 | 0.410 | -0.231 | 0.817 |
| July | -0.211 | 0.423 | -0.498 | 0.619 |
| August | 0.083 | 0.392 | 0.211 | 0.833 |
| September | 0.353 | 0.370 | 0.956 | 0.339 |
| October | 0.470 | 0.361 | 1.303 | 0.192 |
| November | 0.165 | 0.385 | 0.429 | 0.668 |
| December | 0.126 | 0.388 | 0.324 | 0.746 |
| **Habitat × Month** | |  |  |  |
| Center*February | 0.562 | 0.617 | 0.911 | 0.362 |
| Center*March | 0.559 | 0.607 | 0.921 | 0.357 |
| Center*April | 0.656 | 0.601 | 1.092 | 0.275 |
| Center*May | 0.699 | 0.613 | 1.141 | 0.254 |
| Center*June | 0.605 | 0.633 | 0.955 | 0.339 |
| Center*July | 0.513 | 0.657 | 0.781 | 0.435 |
| Center*August | 0.583 | 0.612 | 0.952 | 0.341 |
| Center*September | 0.392 | 0.593 | 0.661 | 0.509 |
| Center*October | 0.117 | 0.597 | 0.197 | 0.844 |
| Center*November | 0.451 | 0.610 | 0.740 | 0.459 |
| Center*December | 0.464 | 0.614 | 0.756 | 0.449 |
| Deep*February | 0.740 | 0.596 | 1.242 | 0.214 |
| Deep*March | 0.529 | 0.598 | 0.884 | 0.376 |
| Deep*April | 0.766 | 0.586 | 1.308 | 0.191 |
| Deep*May | 0.478 | 0.616 | 0.777 | 0.437 |
| Deep*June | 0.456 | 0.633 | 0.721 | 0.471 |
| Deep*July | 0.329 | 0.662 | 0.496 | 0.620 |
| Deep*August | 0.376 | 0.615 | 0.612 | 0.540 |
| Deep*September | 0.752 | 0.565 | 1.331 | 0.183 |
| Deep*October | 0.383 | 0.571 | 0.671 | 0.502 |
| Deep*November | 0.561 | 0.593 | 0.947 | 0.344 |
| Deep*December | 0.715 | 0.589 | 1.213 | 0.225 |

Table A.18. Summary of the GLMM of arthropods species diversity (Simpson diversity, q2) for native species. Species were sampled in the native forest fragment Terra-Brava in three habitats from border (EDGE) to the core forest (DEEP) and in an intermediate habitat (CENTER). Variance explained by fixed effects: Marginal R^2^ = 0.24. Variance explained by the entire model: Conditional R^2^ = 0.24.

|  | ***Estimate*** | ***Std. Error*** | ***z value*** | ***P*** |
| --- | --- | --- | --- | --- |
| **Habitat** |  |  |  |  |
| Intercept | 0.989 | 0.352 | 2.810 | 0.005 |
| Center | 0.415 | 0.454 | 0.914 | 0.361 |
| Deep | 0.183 | 0.477 | 0.385 | 0.700 |
| **Month** |  |  |  |  |
| February | -0.042 | 0.503 | -0.084 | 0.933 |
| March | 0.309 | 0.464 | 0.667 | 0.505 |
| April | 0.109 | 0.485 | 0.225 | 0.822 |
| May | -0.020 | 0.500 | -0.039 | 0.969 |
| June | -0.009 | 0.499 | -0.018 | 0.985 |
| July | 0.783 | 0.425 | 1.842 | 0.066 |
| August | 0.435 | 0.452 | 0.964 | 0.335 |
| September | 0.653 | 0.434 | 1.503 | 0.133 |
| October | 0.210 | 0.474 | 0.444 | 0.657 |
| November | 0.219 | 0.473 | 0.463 | 0.643 |
| December | 0.315 | 0.463 | 0.680 | 0.497 |
| **Habitat × Month** | |  |  |  |
| Center*February | -0.109 | 0.656 | -0.166 | 0.868 |
| Center*March | -1.037 | 0.742 | -1.397 | 0.162 |
| Center*April | -0.584 | 0.670 | -0.872 | 0.383 |
| Center*May | -0.350 | 0.671 | -0.521 | 0.602 |
| Center*June | -0.182 | 0.656 | -0.278 | 0.781 |
| Center*July | -0.348 | 0.562 | -0.619 | 0.536 |
| Center*August | 0.018 | 0.581 | 0.031 | 0.976 |
| Center*September | -0.508 | 0.584 | -0.870 | 0.384 |
| Center*October | -0.752 | 0.669 | -1.125 | 0.260 |
| Center*November | -0.630 | 0.655 | -0.961 | 0.336 |
| Center*December | -0.637 | 0.640 | -0.996 | 0.319 |
| Deep*February | -0.149 | 0.694 | -0.215 | 0.829 |
| Deep*March | -0.679 | 0.684 | -0.993 | 0.321 |
| Deep*April | -0.157 | 0.668 | -0.235 | 0.814 |
| Deep*May | -0.509 | 0.727 | -0.700 | 0.484 |
| Deep*June | -0.217 | 0.694 | -0.313 | 0.754 |
| Deep*July | -0.440 | 0.598 | -0.736 | 0.462 |
| Deep*August | 0.327 | 0.596 | 0.548 | 0.583 |
| Deep*September | -0.284 | 0.603 | -0.472 | 0.637 |
| Deep*October | -0.201 | 0.656 | -0.307 | 0.759 |
| Deep*November | -0.178 | 0.653 | -0.274 | 0.785 |
| Deep*December | -0.550 | 0.669 | -0.822 | 0.411 |

Table A.19. Summary of the GLMM arthropods species diversity (Simpson diversity, q2) for introduced species. Species were sampled in the native forest fragment Terra-Brava in three habitats from border (EDGE) to the core forest (DEEP) and in an intermediate habitat (CENTER). Variance explained by fixed effects: Marginal R^2^ = 0.31. Variance explained by the entire model: Conditional R^2^ = 0.31.

|  | ***Estimate*** | ***Std. Error*** | ***z value*** | ***P*** |
| --- | --- | --- | --- | --- |
| **Habitat** |  |  |  |  |
| Intercept | 3.07E-06 | 7.07E-01 | <0.0001 | 0.999 |
| Center | 5.11E-01 | 8.37E-01 | 0.611 | 0.542 |
| Deep | -2.88E-06 | 9.13E-01 | <0.0001 | 0.999 |
| **Month** |  |  |  |  |
| February | 3.37E-01 | 9.26E-01 | 0.363 | 0.716 |
| March | -7.52E-07 | 1.23E+00 | <0.0001 | 0.999 |
| April | 3.37E-01 | 9.26E-01 | 0.363 | 0.716 |
| May | -2.93E-06 | 1.00E+00 | <0.0001 | 0.999 |
| June | 6.59E-01 | 8.20E-01 | 0.804 | 0.421 |
| July | 1.50E+00 | 7.82E-01 | 1.924 | 0.054 |
| August | 8.28E-01 | 8.04E-01 | 1.031 | 0.303 |
| September | 7.89E-01 | 8.07E-01 | 0.978 | 0.328 |
| October | 1.38E+00 | 7.64E-01 | 1.808 | 0.071 |
| November | 1.28E+00 | 7.70E-01 | 1.655 | 0.098 |
| December | 7.89E-01 | 8.07E-01 | 0.977 | 0.329 |
| **Habitat × Month** | |  |  |  |
| Center*February | -8.47E-01 | 1.25E+00 | -0.679 | 0.497 |
| Center*March | 7.62E-01 | 1.41E+00 | 0.541 | 0.588 |
| Center*April | -8.47E-01 | 1.43E+00 | -0.591 | 0.555 |
| Center*May | -5.11E-01 | 1.30E+00 | -0.392 | 0.695 |
| Center*June | -4.77E-01 | 1.17E+00 | -0.407 | 0.684 |
| Center*July | -1.32E+00 | 9.89E-01 | -1.337 | 0.181 |
| Center*August | -1.28E+00 | 1.08E+00 | -1.189 | 0.234 |
| Center*September | -5.18E-01 | 1.00E+00 | -0.517 | 0.605 |
| Center*October | -1.12E+00 | 9.68E-01 | -1.152 | 0.249 |
| Center*November | -9.02E-01 | 9.65E-01 | -0.934 | 0.350 |
| Center*December | -9.46E-01 | 1.04E+00 | -0.908 | 0.364 |
| Deep*February | -3.37E-01 | 1.30E+00 | -0.259 | 0.796 |
| Deep*March | -4.69E-07 | 1.68E+00 | <0.0001 | 0.999 |
| Deep*April | 5.80E-01 | 1.18E+00 | 0.492 | 0.623 |
| Deep*May | 6.93E-01 | 1.23E+00 | 0.566 | 0.571 |
| Deep*June | -3.72E-01 | 1.12E+00 | -0.332 | 0.740 |
| Deep*July | -2.71E-01 | 1.02E+00 | -0.265 | 0.791 |
| Deep*August | 6.54E-01 | 1.03E+00 | 0.636 | 0.525 |
| Deep*September | 3.20E-01 | 1.05E+00 | 0.306 | 0.760 |
| Deep*October | -8.97E-02 | 1.00E+00 | -0.089 | 0.929 |
| Deep*November | 1.92E-01 | 1.00E+00 | 0.191 | 0.848 |
| Deep*December | 2.55E-01 | 1.05E+00 | 0.242 | 0.809 |
